# TRPM7 kinase-mediated immunomodulation in macrophage plays a central role in magnesium ion-induced bone regeneration

**DOI:** 10.1101/2020.04.24.059881

**Authors:** Wei Qiao, Karen H.M. Wong, Jie Shen, Wenhao Wang, Jun Wu, Jinhua Li, Zhengjie Lin, Zetao Chen, Jukka P. Matinlinna, Yufeng Zheng, Shuilin Wu, Xuanyong Liu, Keng Po Lai, Zhuofan Chen, Yun Wah Lam, Kenneth M.C. Cheung, Kelvin W.K. Yeung

**Affiliations:** Department of Orthopaedics and Traumatology, Li Ka Shing Faculty of Medicine, the University of Hong Kong, Hong Kong S.A.R., P.R. China; Shenzhen Key Laboratory for Innovative Technology in Orthopaedic Trauma, The University of Hong Kong-Shenzhen Hospital, Shenzhen 518053, P.R. China; Dental Materials Science, Applied Oral Sciences, Faculty of Dentistry, the University of Hong Kong, Hong Kong S.A.R., P.R. China; Department of Oral Implantology, Hospital of Stomatology, Guanghua School of Stomatology, Institute of Stomatological Research, Sun Yat-sen University, Guangzhou 510030, P. R. China; Zhujiang New Town Clinic, Hospital of Stomatology, Sun Yat-sen University, Guangzhou, 510000, P. R. China; Centre for Translational Bone, Joint and Soft Tissue Research, University Hospital and Faculty of Medicine Carl Gustav Carus, Technische Universität Dresden, Dresden, 01307, Germany; Department of Chemistry, City University of Hong Kong, Kowloon Tong, Hong Kong S.A.R., P.R. China; State Key Laboratory for Turbulence and Complex System and Department of Materials Science and Engineering, College of Engineering, Peking University, Beijing 100871, P.R. China; School of Materials Science & Engineering, Tianjin University, Tianjin 300072, P.R. China; State Key Laboratory of High Performance Ceramics and Superfine Microstructure, Shanghai Institute of Ceramics, Chinese Academy of Sciences, Shanghai 200050, P.R. China; China Orthopedic Regenerative Medicine Group (CORMed), Hangzhou 310000, P.R. China

**Author notes:** These authors share the corresponding authorship. **Dr. Kelvin W.K. Yeung**, Department of Orthopaedics and Traumatology, Li Ka Shing Faculty of Medicine, the University of Hong Kong, Hong Kong S.A.R., P.R. China., **Dr. Yun Wah Lam**, Department of Chemistry, City University of Hong Kong, Tat Chee Road, Kowloon Tong, Hong Kong S.A.R., P.R. China., **Prof. Zhuofan Chen**, Department of Oral Implantology, 5F, Hospital of Stomatology, Sun Yat-sen University, 56 Ling Yuan Xi Road, Guangzhou, 510030, P. R. China.

**Keywords:** Magnesium (Mg), bone regeneration, Transient receptor potential cation channel member 7 (TRPM7), macrophage

## Abstract

The use of magnesium ion (Mg^2+^)-modified biomaterials in bone regeneration is a promising and cost-effective therapeutic. Despite the widespread observation on the osteogenic effects of Mg^2+^, the diverse roles played by Mg^2+^ in the complex biological process of bone healing have not been systematically dissected. Here, we reveal a previously unknown biphasic mode of action of Mg^2+^ in bone repair. In the early inflammation phase, Mg^2+^ primarily targets the monocyte-macrophage lineage to promote their recruitment, activation, and polarization. We showed that an increase in extracellular Mg^2+^ contributes to an upregulated expression of transient receptor potential cation channel member 7 (TRPM7) and a TRPM7-dependent influx of Mg^2+^ in the monocyte-macrophage lineage, resulting in the cleavage and nuclear accumulation of TRPM7-cleaved kinase fragments (M7CKs). This then triggers the phosphorylation of Histone H3 at serine 10, in a TRPM7-dependent manner at the promoters of inflammatory cytokines like IL-8, leading to the formation of a pro-osteogenic immune microenvironment. In the later active repair/remodeling phase of bone healing, however, continued exposure of Mg^2+^ and IL-8 leads to over activation of NF-κB signaling in macrophages, turning the immune microenvironment into pro-osteoclastogenesis. Moreover, the presence of Mg^2+^ at this stage also decelerates bone maturation through the suppression of hydroxyapatite precipitation. The negative effects of Mg^2+^ on osteogenesis can override the initial pro-osteogenic benefits of Mg^2+^, as we found prolonged delivery of Mg^2+^ compromises overall bone formation. Taken together, this study establishes a paradigm shift in understanding the diverse and multifaceted roles of Mg^2+^ in bone healing.

## Introduction

Bone tissue has a substantial capacity for repair and regeneration after injury or surgical treatment. However, the natural healing of bone can be a slow process and often fails to restore the bone to its original strength and structure ^1^. Thus, clinical interventions using orthopedic biomaterials are often required to accelerate bone healing while maintaining the amount and quality of bone mass. As one of the essential ions in metabolism, magnesium ion (Mg^2+^) is integral to bone homeostasis and metabolism. Deficiency in magnesium is known to disrupt systemic bone metabolism, characterized by inadequate bone formation and deregulated bone resorption ^2–8^. In contrast, Mg^2+^ supplement is beneficial to patients of osteoporosis ^9^. Magnesium, its alloys ^10–13^ and derivatives ^14–16^ have been extensively studied as replacements for non-degradable metallic implants e.g., titanium alloys in bone surgeries. Mg^2+^ modified biomaterials have shown superior osteogenic capacity in many reports ^14–21^. However, the detrimental effects of Mg^2+^ released upon degradation have also been observed ^22–24^. These conflicting results may reflect the incomplete understanding of the roles of Mg^2+^ in the complex biological process of bone healing.

Our group has recently reported that the incorporation of Mg^2+^ in polycaprolactone (PCL) implant ^18^ and poly(lactic-co-glycolic acid) (PLGA) microsphere ^19^ can promote bone formation in a rat femoral defect model. We identified ~50-200 ppm as the optimal Mg^2+^ concentration for promoting the osteogenic activities of osteoblast *in vitro*, as well as new bone formation *in vivo* ^13,18^. Moreover, by using customized biomaterials that enable the controlled release of Mg^2+^ at different stages of bone healing, we demonstrated that the bone regeneration rate and the quality of newly formed bone tissues depend on the release profile of Mg^2+^ ^18,19^. Bone healing is a complex process that involves the precise coordination of osteoclastogenesis and osteogenesis, through the interplay of multiple types of cells in a dynamic microenvironment. It is possible that the different cell types involved in various phases of bone healing, from early inflammation to the later bone formation and remodelling, may respond to Mg^2+^ in different ways.

The monocyte-macrophage cell lineage has been recognized as a major player in acute inflammation response to biomaterials, mainly due to their high plasticity in response to environmental cues and their multiple roles in the bone homeostasis. According to their distinct functional properties, surface markers, and inducer, macrophages are characterized into several phenotypes (*i.e.*, M1, M2a, M2b, and M2c) ^25^. The doping of Mg^2+^ into titanium ^26^ and calcium phosphate cement ^27^ were demonstrated to promote the M2 polarization of macrophages, resulting in the decrease of pro-inflammatory cytokines and the increase of anti-inflammatory cytokines. In fact, magnesium has been characterized as an anti-inflammatory agent that can reduce the expression and release of pro-inflammatory molecules. Interestingly, Mg^2+^ deficiency can increase the production of cytokines that contribute to osteoclastogenesis ^28,29^, whereas Mg^2+^ supplement suppresses the secretion of pro-inflammatory cytokines and upregulates tissue repair factors ^30,31^. However, there is yet no consensus as to which macrophage phenotype is more beneficial to bone regeneration, because both M1 ^32,33^ and M2 ^34,35^ phenotypes have been reported to contribute to osteogenesis in specific ways. Moreover, the conventional M1/M2 classification of macrophages has recently been challenged by a more heterogeneous grouping method, which suggests there may exist a continuum between M1 and M2 phenotypes yet to be identified ^36^. Thus, the complexity of the Mg^2+^-induced immunomodulation on macrophages as well as its specific effects on the bone healing process in the complicated *in vivo* scenario requires further investigation.

In the present study, we systematically analyzed the dose- and time-dependent effects of Mg^2+^ on monocyte-macrophage lineage and osteoblast lineage involved in bone healing and investigated the underlying mechanisms behind the action of Mg^2+^. We demonstrated a previously undefined immunomodulatory role of Mg^2+^ at the early phase of bone healing, in which macrophages are stimulated, through a signalling pathway that involves transient receptor potential cation channel member 7 (TRPM7), to generate a specific pro-osteogenic immune microenvironment. At later bone repair/remodeling phase, the presence of Mg^2+^ impacts on bone healing in another way, through activation of NF-κB signaling-mediated osteoclastogenesis and inhibition on mineralization of extracellular matrix. We believe that these results will inspire the development of next generation of Mg^2+^-based degradable biomaterials for clinical use that can better harness the healing power of Mg^2+^.

## Results

### The time-dependent effect of Mg^2+^ on bone regeneration

To elucidate the time-dependent effect of Mg^2+^ on bone healing, we developed an alginate-based hydrogel that allows a transient release of MgCl_2_, at a concentration of ~10 mM, over one week (Fig. S1a). Critical-sized tunnel defects with a diameter of 2 mm were created in the distal end of rat femora ^18,19^. Mg^2+^ hydrogel was injected into the cavities at different time points after the injury, causing a temporary and localized increase in Mg^2+^ to around 10x of the physiological level in the defect. The impact of this implant on chloride levels is negligible, as the physiological chloride concentration is over 100 times higher than Mg^2+^ concentration ^37^. Using scanning electron microscopy with energy dispersive X-Ray spectroscopy (SEM-EDX), we demonstrated that the Mg^2+^ releasing hydrogel significantly increased the magnesium content while decreasing the calcium content in the defects on day 7 post-injury (Fig. S1c-e). Animals were randomly assigned into one of three treatment regimens; (1) injection of Mg^2+^ hydrogel directly after injury (Fig. 1a); (2) injection at day 7 post-injury (Fig. 2a) and (3) injection at both day 0 and day 7 post-injury (Fig. 2g). Controls for each treatment regimen were animals injected with an equivalent dose of alginate hydrogel without Mg^2+^. The efficiency of bone repair of these animals was monitored over 8 weeks, compared to a sham group in which the injured animals were not injected with any hydrogel. Micro-CT analyses showed that no observable healing in the sham group after 8 weeks (Fig. 1b, S1f). Transient exposure of Mg^2+^ during the first week after injury *(Regimen 1)* led to a 3-fold increase in the trabecular bone fraction (BV/TV), a 2.5-fold increase in trabecular number (Tb. N), a 2-fold increase in bone mineral density (BMD of TV and BMD of BV) and a 2-fold increase in trabecular thickness (Tb.Th), as compared to the control and the sham groups (Fig. 1b, c, S1f, g). Nanoindentation test showed Mg^2+^ hydrogel promoted new bone formation without compromising the mechanical properties, as Young’s modulus of the Mg^2+^-induced newly formed bone at post-operation week 8 was comparable to the control and sham group (Fig. S1b). Histological assessments by H&E staining (Fig. 1d, S1h) confirmed the increase in bone formation in the defects grafted with Mg^2+^ releasing hydrogel. Meanwhile, the number of osteocalcin (OCN) positive osteoblasts was increased in the Mg^2+^ treated group compared to the control and sham groups (Fig. 1e, g, S1j); whereas, the number of tartaric acidic phosphatase (TRAP) positive osteoclasts in the Mg^2+^ treated group was the lowest (Fig. 1f, g). Calcein/Xylenol labeling (Fig. 1i, S1i) and Goldner’s trichrome staining (Fig 1j) demonstrated that new bone formation and mineral apposition were more active in the Mg^2+^ treated group than in the control.

**Fig. 1:**
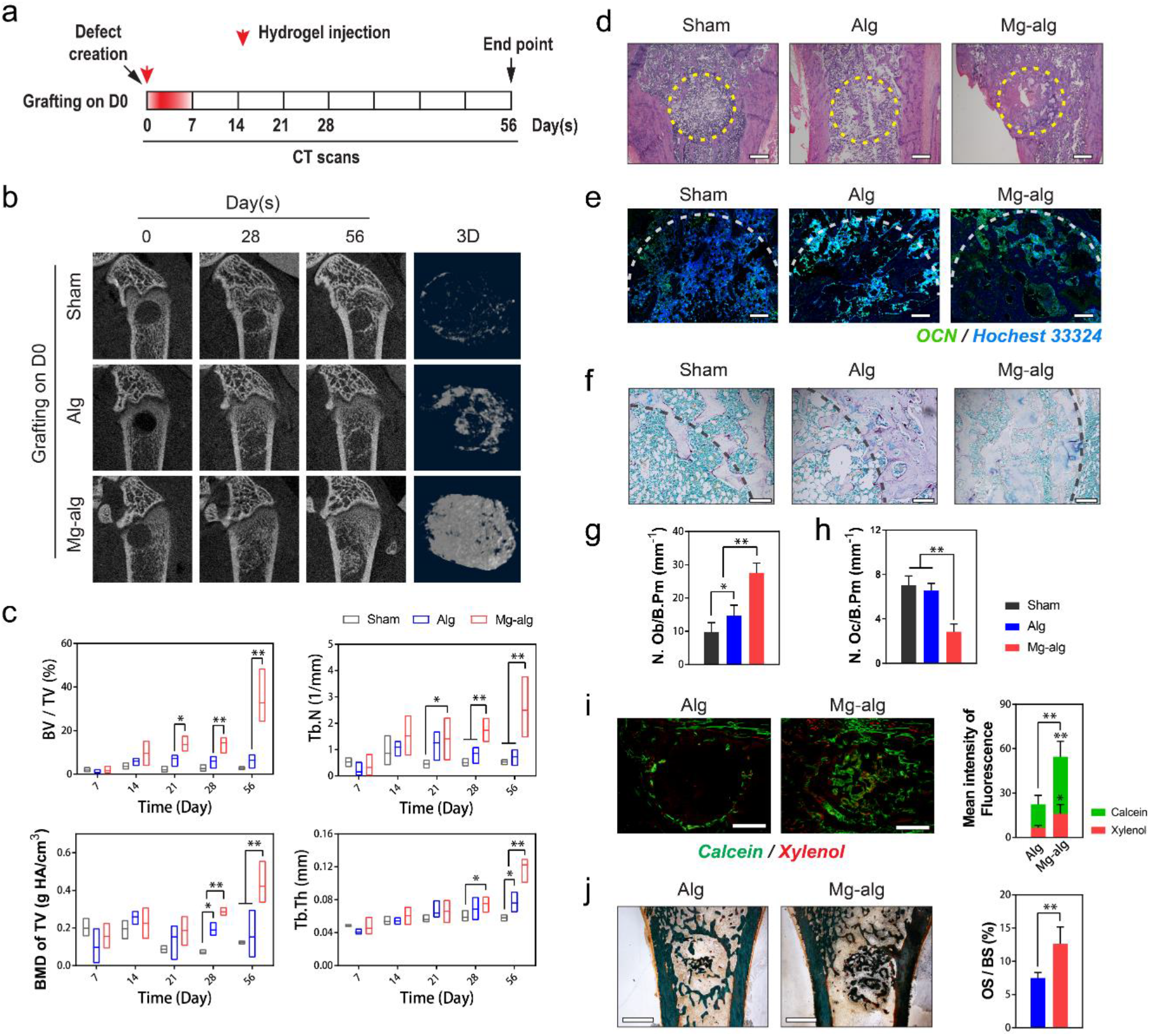
Effects of Mg^2+^ releasing alginate on the bone healing of defects in rat femur. **(a)** Mg^2+^-crosslinked alginate was injected into the femur defect in rats right after the injury, hence the release of Mg^2+^ was limited to the first week of injury. **(b)** Representative micro-CT images and reconstructed 3D images of the defects in rat femora without grafting (Sham group, n = 3), grafted with pure alginate (serve as a control, n = 5) or Mg^2+^ releasing alginate (n = 6). **(c)** Corresponding measurements of trabecular bone fraction (BV/TV), trabecular number (Tb.N), bone mineral density (BMD of TV) and trabecular thickness (Tb.Th) showing the healing process of rat femoral defects from day 7 to day 56. \**P*<0.05, \*\**P*<0.01 by one-way ANOVA with Tukey’s *post hoc* test. **(d)** Representative H&E staining images of the grafted defects in the rat femora, scale bars =500 μm. **(e)** Representative immunofluorescent images showing the expression of OCN in the grafted defects in the rat femora, scale bars =500 μm. **(f)** Representative TRAP staining images showing the presence of osteoclasts in the grafted defects in the rat femora, scale bars =200 μm. **(g,h)** Histomorphological analysis of osteoblast numbers (**g**, N.Ob/B.Pm) and osteoclast numbers (**h**, N.Oc/B.Pm) in the defects of femora. **(i)** Representative images of calcein/xylenol labeling for bone regeneration in the rat femoral defects grafted with alginate or Mg^2+^ releasing alginate, scale bars = 1 mm. **(j)** Representative Goldner’s trichrome staining of the grafted defects on day 56, scale bars = 1 mm, and quantitative analysis of osteoid surface per bone surface (OS/BS) in the grafted femoral defects.

**Fig. 2.**
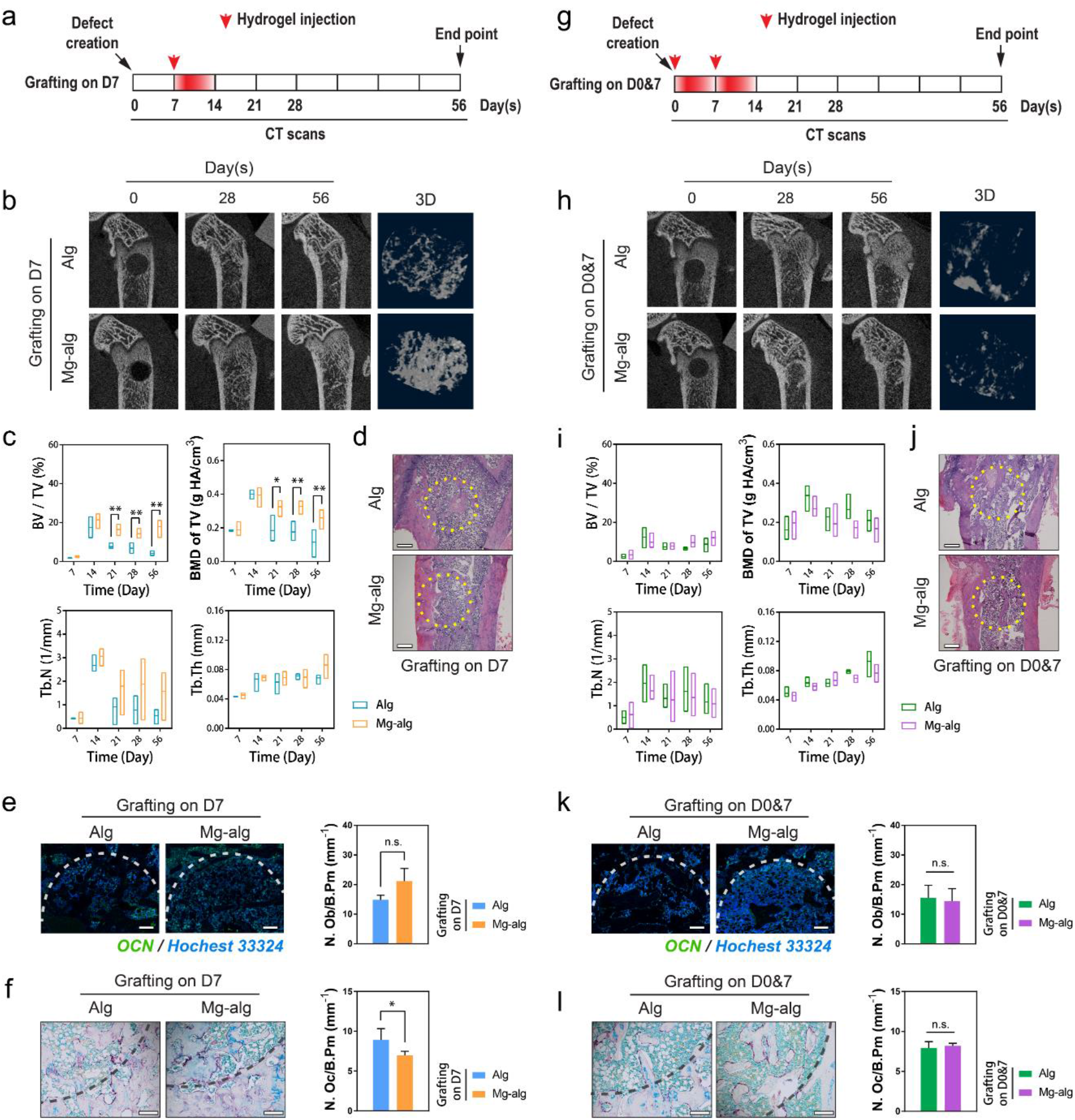
**(a)** Mg^2+^-crosslinked alginate was injected into the femur defect in rats at the seventh days after the injury to exclude the effects of Mg^2+^ on early phrase inflammation. **(b)** Representative micro-CT and reconstructed 3D images of the defects in rat femur on day 56 when the grafting was delayed. **(c)** Corresponding measurements of trabecular bone fraction (BV/TV), trabecular number (Tb.N), bone mineral density (BMD of TV) and trabecular number (Tb.Th) showing the healing process of rat femoral defects from day 7 to day 56. \**P*<0.05, \*\**P*<0.01 by one-way ANOVA with Tukey’s *post hoc* test. **(d)** Representative H&E staining images of the grafted defects in the rat femora, scale bars =500 μm. **(e)** Representative immunofluorescent images showing the expression of OCN in the grafted defects in the rat femora, scale bars =500 μm. **(f)** Representative TRAP staining images showing the presence of osteoclasts in the grafted defects in the rat femora, scale bars =200 μm. **(g)** Mg-crosslinked alginate was injected into the femur defect in rats at both the first and seventh days after the injury to allow sustained release of Mg^2+^ in the first two weeks of injury. **(h)** Representative micro-CT and reconstructed 3D images of the defects in rat femur on day 56 when the grafting was repeated. **(i)** Corresponding measurements of trabecular bone fraction (BV/TV), trabecular number (Tb.N), bone mineral density (BMD of TV) and trabecular number (Tb.Th) showing the healing process of rat femoral from day 7 to day 56. **(j)** Representative H&E staining images of the grafted defects in the rat femora, scale bars =500 μm. **(k)** Representative immunofluorescent images showing the expression of OCN in the grafted defects in the rat femora, scale bars =500 μm. **(l)** Representative TRAP staining images showing the presence of osteoclasts in the grafted defects in the rat femora, scale bars =200 μm.

Despite a higher trabecular bone fraction and bone mineral density, the overall beneficial effects of Mg^2+^ were significantly attenuated when the delivery of the Mg^2+^, even at the same dose, was delayed to the second week (*Regimen 2*, Fig. 2b, c, S2a, b). There was no significant difference in the bone morphology according to H&E staining (Fig. 2d, S2c). Indeed, the number of TRAP+ osteoclasts was only slightly lower when the Mg^2+^ releasing hydrogel was applied at the second week of bone healing (Fig. 2f), while the number of OCN+ osteoblasts remains unchanged (Fig. 2e, S2d). When Mg^2+^ was continuously delivered over the first two weeks post-injury *(Regimen 3*, Fig. 2g), the benefit of Mg^2+^ hydrogel on bone formation measured by micro-CT became negligible (Fig. 2h, i, Fig. S2e, f). In addition, the bone morphology, as well as the number of OCN+ osteoblasts and TRAP+ osteoclasts were not different between Mg^2+^ treated group and the control (Fig. 2j-l, S2g, h). These observations demonstrate that the Mg^2+^ promotes bone healing only when delivered during the initial phase of repair, and a prolonged treatment resulted in unexpected detrimental effects on bone formation.

### Mg^2+^-induced bone regeneration is mediated by macrophage activities

As the effective window of Mg^2+^ exposure coincided with the initial phase of inflammation dominated by macrophages ^38^, it is possible that the effect of Mg^2+^ on bone formation might be mediated through its modulation of macrophages. We first demonstrated that the use of Mg^2+^ releasing hydrogel contributed to a significant increase in the number of CD68 positive macrophages in the proximity of the defect on day 7 after the operation (Fig. 3b and Fig. S3b, c). However, the effect of Mg^2+^ on the recruitment of macrophages vanished when the macrophage activity of rats was selectively depleted in the first week after the surgery by intraperitoneal administrations of liposome-encapsulated clodronate ^39^ (Fig. 3a, b). Meanwhile, the addition of Mg^2+^ also contributed to a group of macrophage-derived TRAP+ preosteoclasts at the early stage (i.e., day 7) of bone healing, which disappeared in the macrophage-depleted model (Fig. 3c). Compared with the vehicle group showing the release of Mg^2+^ promotes new bone formation, the osteopromotive effects of Mg^2+^ in macrophage-depleted rats were reversed even when it is delivered in the optimal time window. Indeed, Mg^2+^ exposure appeared to delay bone healing in these animals: the BV/TV was lower in Mg^2+^ treated group from day 14 to 28 relative to the control group (Fig 3d, f, S3a), while the histological analysis demonstrated that the effects of Mg^2+^ on promoting osteogenesis while suppressing osteoclastogenesis became insignificant (Fig. 3e, g, h, S3b, d).

**Fig. 3:**
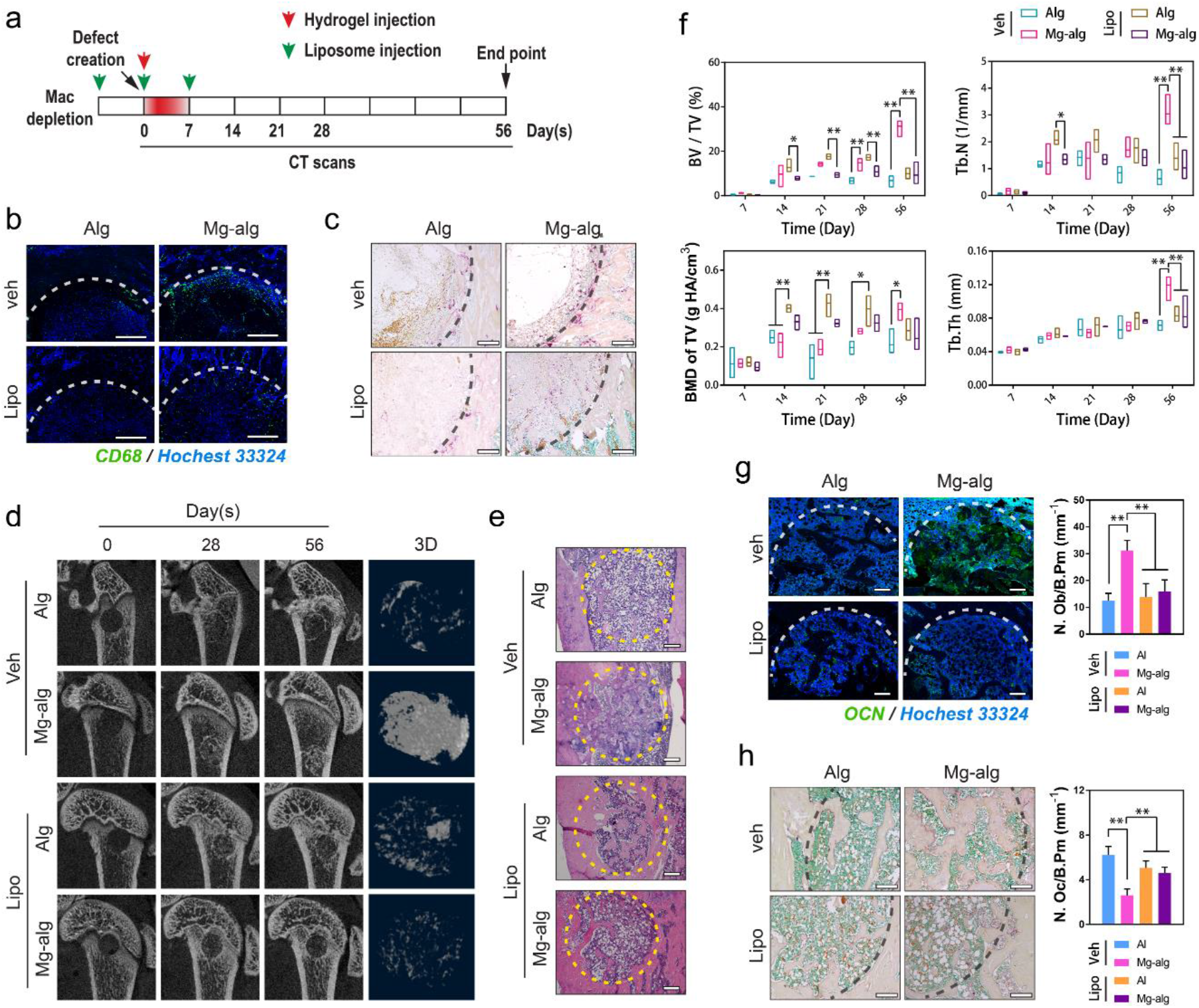
Mg^2+^-induced immunomodulation of macrophages. **(a)** Mg-crosslinked alginate was injected into the femur defect in rats when their macrophages were selectively depleted by intraperitoneal administrations of liposome-encapsulated clodronate. **(b)** Representative immunofluorescent images showing the infiltration of CD68+ macrophages on day 7 in the grafted defects in the rat femora, scale bars =500 μm. **(c)** Representative TRAP staining images showing the presence of osteoclasts in the grafted defects in the rat femora on day 7, scale bars =200 μm. **(d)** Representative micro-CT and reconstructed 3D images of the defects in rat femur on day 56 when the grafting was delayed. **(e)** Representative H&E staining images of the grafted defects in the rat femora, scale bars =500 μm. **(f)** Corresponding measurements of trabecular bone fraction (BV/TV), trabecular number (Tb.N), bone mineral density (BMD of TV) and trabecular number (Tb.Th) showing the healing process of rat femoral defects from day 7 to day 56. \**P*<<0.05, \*\**P*<0.01 by one-way ANOVA with Tukey’s *post hoc* test. **(g)** Representative immunofluorescent images showing the expression of OCN in the grafted defects in the rat femora, scale bars =500 μm. **(h)** Representative TRAP staining images showing the presence of osteoclasts in the grafted defects in the rat femora, scale bars =200 μm.

We conducted *in vitro* experiments to further delineate the role of Mg^2+^ in macrophage functions by exposing THP1, a human monocyte cell line that can be differentiated into macrophages, to different Mg^2+^ concentrations (named 0.1x, 1x and 10x to represent the Mg^2+^ concentrations in each medium relative to that in a physiological Mg^2+^ level in the culture medium, 0.8 mM). An increase of Mg^2+^ concentration in the culture medium significantly promoted the maturation of suspended monocytes into adhered macrophages (Fig. 4a, S4c). Moreover, Mg^2+^ also increased the activity of THP1-derived macrophages, as evidenced by the increase of intracellular ATP levels (Fig. 4b) and the number of mitochondria (Fig. S4e). Furthermore, the presence of Mg^2+^ protected the macrophage from ROS production under Mg^2+^ insufficient condition (Fig. 4c). Immunophenotyping using RT-qPCR demonstrated that 10x Mg^2+^ treatment resulted in increased expression of M2 macrophage surface markers CD163 and CD206 (Fig. 4d). This is further supported by flow cytometry data showed that the number of macrophages expressing M2 surface markers CD163 and CD206 was increased by the stimulation of 10x Mg^2+^, however, the number of M1 macrophages characterized by the expression of CD80 remained unchanged (Fig. 4e). Our RT-qPCR data also revealed the effect of 10x Mg^2+^ on upregulating a series of genes encoding cytokines favoring osteogenesis, such as CCL5, IL-1ra, IL-8, TGF-β1, BMP2, VEGFA, IL-10 (Fig. 4f), while downregulating genes encoding cytokines favoring osteoclastogenesis, including OSM, IL-6, IL-1β, TNF-α (Fig. 4g). Using cytokine array, the major cytokines secreted by macrophages upon the stimulation of Mg^2+^ were found to be IL-1ra, IL-8 and CCL5, which were distinct from traditionally characterized M1 or M2 phenotypes (Fig. 4h, S4g). We then confirmed the effects of Mg^2+^ on increasing the expression of IL-1ra, IL-8 and CCL5 while suppressing the expression of IL-1β by western blots (Fig. 4i). Our *in vitro* findings were supported by our immunostaining and ELISA data *in vivo* showing that Mg^2+^ released from the hydrogel led to a significant increase in the level of IL-8 (Fig. 3l, m, S3c), CCL5 (Fig. S4a) and IL-1ra (Fig. S4b), as well as a significant decrease in the level of IL-1β (Fig.3n, o, S3c) in the femoral defects one week post-operation. However, in macrophage-depleted animals, the level of these cytokines remained low and unchanged regardless of the addition of Mg^2+^ in the hydrogel. We also demonstrated that the effects of Mg^2+^ on the levels of IL-8 and IL-1β to be both time-dependent and concentration-dependent using THP1-derived macrophages (Fig. 4j, k, p, q).

**Fig. 4:**
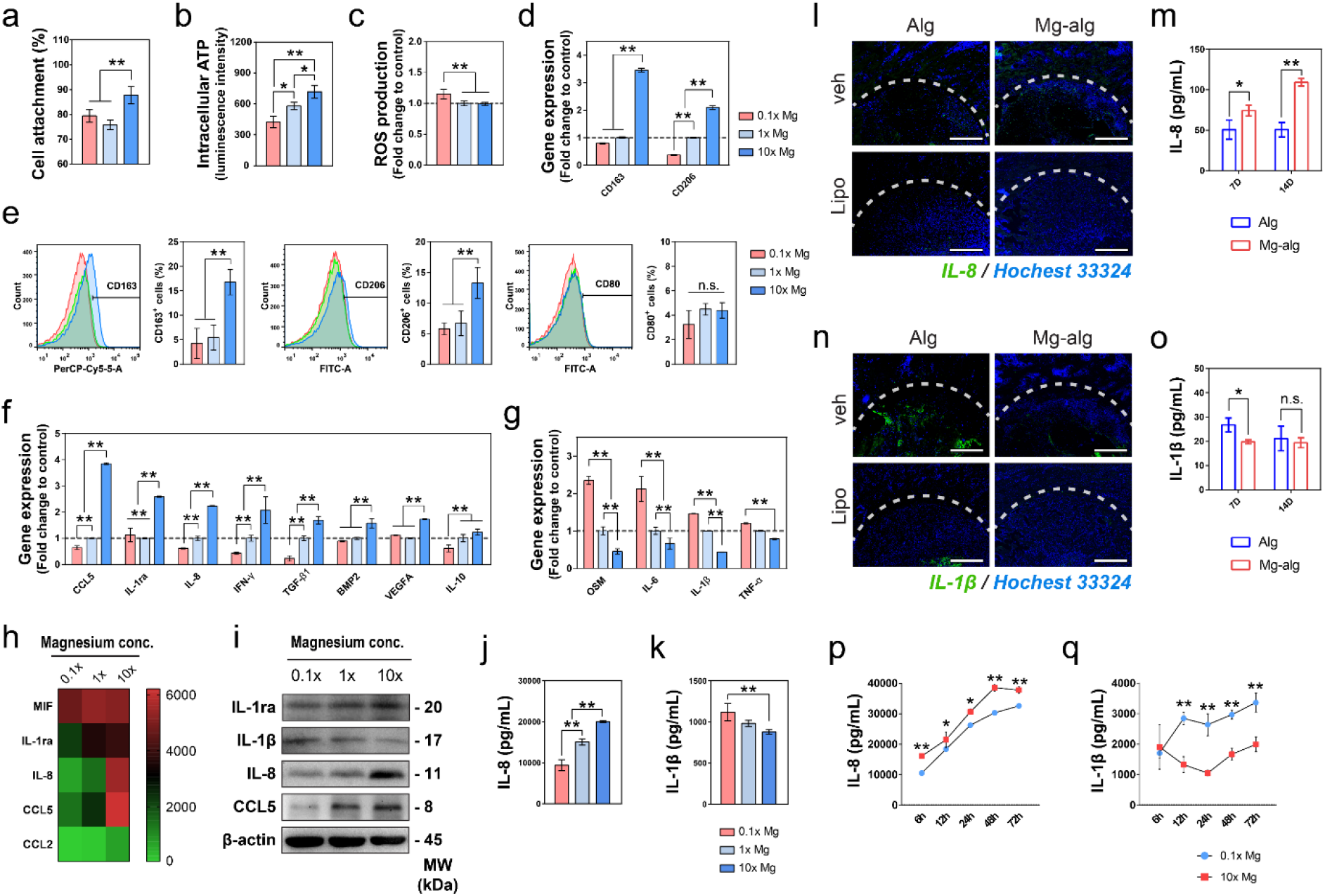
Mg^2+^-induced immunomodulation of macrophages. **(a, b, c)** The effects of different concentrations of Mg^2+^ on the cell attachment (**a**, n ≥ 5), intracellular ATP level (**b**, n ≥ 5) and ROS production (**c**, n ≥ 5) of macrophages differentiated from suspended THP1 monocytes. The data for cell attachment was expressed as a percentage of initially seeded THP-1 cells. \**P*<0.05, \*\**P*<0.01 by one-way ANOVA with Tukey’s *post hoc* test. **(d)** The effect of different concentrations of Mg^2+^ on the gene expression of CD163 and CD206 in macrophages evaluated by RT-qPCR. **(e)** The effect of different concentrations of Mg^2+^ on the polarization of macrophages was evaluated by the expression of CD163, CD206, and CD80 using flow cytometry. **(f, g)** The inflammatory-related genes upregulated **(f)** and downregulated **(g)** by the stimulation of Mg^2+^. Data are mean ± s.d. \**P*<0.05, \*\**P*<0.01 by one-way ANOVA with Tukey’s *post hoc* test. **(h)** Major cytokines that respond to the stimulation of Mg^2+^ determined by cytokine arrays were shown in a heat map. **(i)** Representative western blots showing the expression of IL-1ra, IL-1β, IL-8, and CCL5 of macrophages cultured in medium supplemented with different concentrations of Mg^2+^. **(j,k)** ELISA analysis showing the concentration-dependent effect of Mg^2+^ on the production of IL-8 **(j)** and IL-1β **(k)** in THP1-derived macrophages. **(l, m)** Representative immunofluorescent images **(l)** and Elisa analysis **(m)** showing the expression of IL-8 on day 7 in the grafted defects in the rat femora, scale bars =500 μm. **(n, o)** Representative immunofluorescent images **(n)** and Elisa analysis **(o)** showing the expression of IL-1β on day 7 in the grafted defects in the rat femora, scale bars =500 μm. **(p,q)** ELISA analysis showing the time-dependent effect of Mg^2+^ on the production of IL-8 **(p)** and IL-1β **(q)** in THP1-derived macrophages.

### The central role of TRPM7 in Mg^2+^-induced inflammatory modulation in macrophages

Using Mag-Fluo-4, an Mg^2+^ specific dye, we observed a rapid increase of intracellular Mg^2+^ levels in macrophage upon the addition of Mg^2+^ in the culture medium, peaking at ~8 min before reaching a steady state higher than the baseline (Fig. 5a, b, S4f). Our ICP-OES data also verified that the Mg^2+^ level was consistently higher than the baseline when cultured in supplemented medium (Fig. S4d). As expected, when the channel activity of TRPM7, a Mg^2+^ transporter, was inhibited by FTY720, a potent TRPM7 blocker that inhibits channel activity by reducing the open probability ^40^, the inflow of Mg^2+^ became insignificant (Fig. 5a). In addition, our RT-qPCR data showed that 10x Mg^2+^ contributed to a more than two-fold upregulation in the TRPM7 gene relative to the marginal increase in the MagT1 gene (Fig.5c). Using an antibody that targets the C-terminal of TRPM7, a region containing the TRPM7-cleaved kinase fragments (M7CKs), we demonstrated that Mg^2+^ treatment increased the overall level of full-length TRPM7 in macrophages, with a corresponding increase in M7CKs (Fig. 5e). This was also validated *in vivo* by increased expression of TRPM7 at the proximity of the Mg-alg grafted defect compared with Alg grafted control (Fig. S4h, i). Immunofluorescence staining (Fig. 5d) and western blots analysis of subcellular fractionation (Fig. 5f) confirmed the accumulation of M7CKs in macrophage nuclei upon the stimulation of Mg^2+^. Moreover, the phosphorylation of Histone H3 at residue S10 (H3S10p) was found to be increased in macrophages after Mg^2+^ treatment (Fig. 5f, Fig. S4j), consistent with the Mg^2+^ induced accumulation of M7CKs. Using super-resolution microscopy, we demonstrated that nuclear M7CKs localized in foci that overlapped with a subset of H3 foci (Fig. S4k). Therefore, we performed chromatin immunoprecipitation (ChIP) assays and found that Mg^2+^ increased the association with the IL-8 promoter of Histone H3 that was phosphorylated on Ser 10 (Fig. 5g). Moreover, the gene loci at the proximity of IκBα promoter was also associated with pH3S10-containing chromatin in an Mg^2+^-dependent manner (Fig. 6h).

**Fig. 5:**
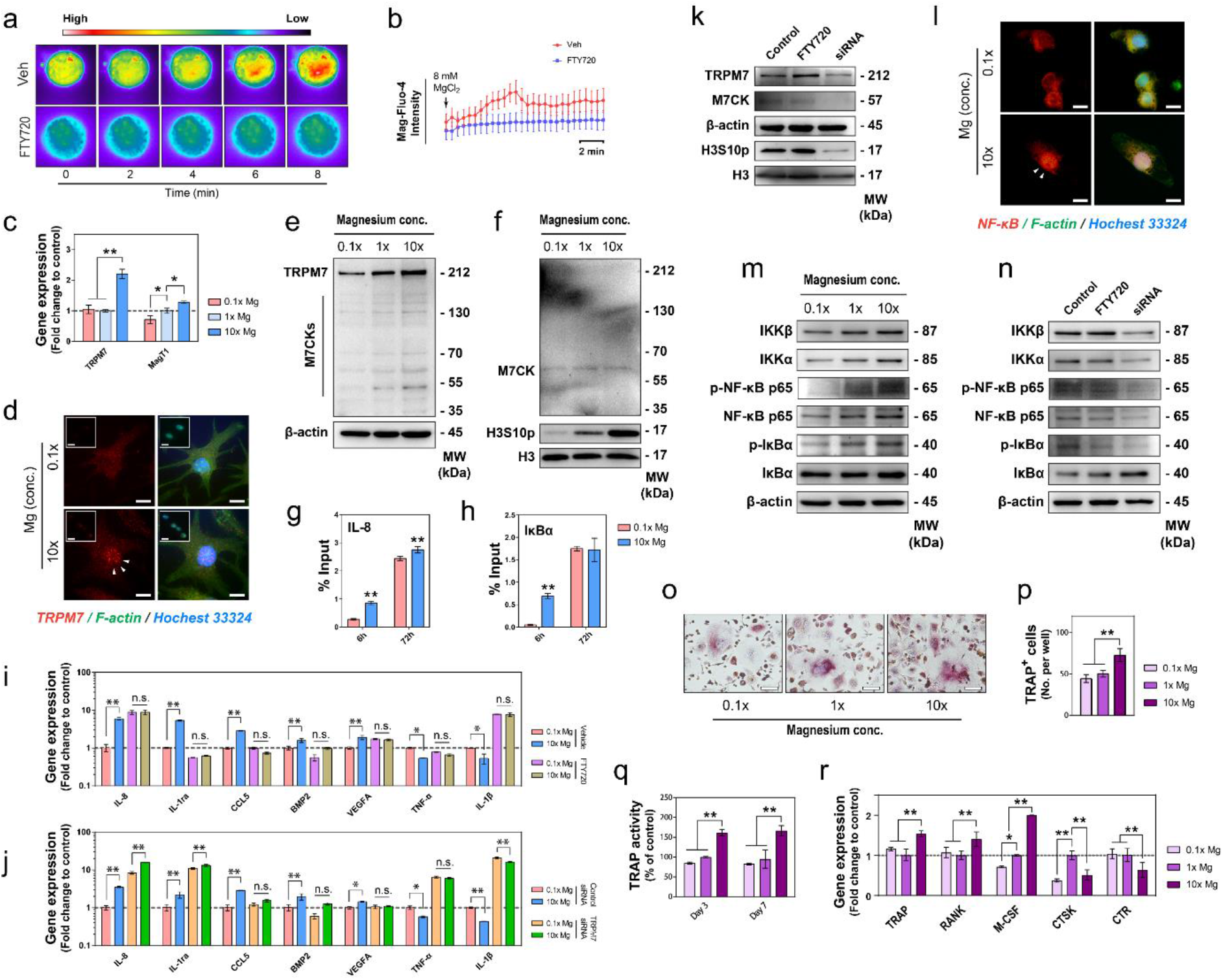
The involvement of TRPM7 in Mg-induced immunomodulation in macrophages. **(a,b)** Representative fluorescence images **(a)** showing the entry of Mg^2+^ into macrophages after the addition of 8 mM MgCl_2_. Color scale bar from low (black to blue) to high (red to white) indicates the level of Mg^2+^. **(b)** Time-course changes in intracellular Mg^2+^ quantified by measuring the intensity of fluorescence. Scale bar = 2 min. **(c)** The gene expression of TRPM7 and MagT1 in THP1-derived macrophages upon the stimulation of different concentrations of Mg^2+^ determined by RT-qPCR. **(d)** Representative fluorescence images showing the nucleus accumulation of TRPM7 in macrophages after the stimulation of Mg^2+^. Inserts showed the staining with the cell permeabilization before the fixation to better demonstrate the nucleus bound TRPM7. Scale bars = 5 μm. **(e)** Representative western blots showing the effects of Mg^2+^ on the expression of TRPM7 and its cleaved kinase fragments (M7CKs) in THP-1 derived macrophages. **(f)** Nuclear protein probed with western blots showing the stimulation of Mg^2+^ contributed to increased nuclear fraction of M7CK at around 60 kDa. Meanwhile, the phosphorylation of Histone H3S10 was upregulated by the addition of Mg^2+^. **(g,h)** The phosphorylation of Histone H3S10 at promoters of IκBα **(g)** and IL-8 **(h)** was upregulated by the stimulation of Mg^2+^. **(i,j)** The effects of TRPM7 siRNA **(i)** and FTY720 **(j)** on the inflammatory and bone-related gene expression in macrophages. Data are mean ± s.d. \**P*<0.05, \*\**P*<0.01 by one-way ANOVA with Tukey’s *post hoc* test. **(k)** Representative western blots showing the effects of FTY720 or siRNA on the expression of TRPM7, M7CK, and the phosphorylation of Histone H3S10. **(l)** Representative fluorescence images showing the nucleus translocation of NF-κB in macrophages after the stimulation of Mg^2+^. **(m, n)** Representative blots showing the concentration-dependent effect of Mg **(m)** and the influence of FTY720 or siRNA **(n)** on the activation of NF-κB signaling. **(o,p)** Representative microscopic images **(o)** and quantitative data **(p)** showing the formation of multi-nuclear TRAP+ cells from macrophages stimulated with different concentrations of Mg^2+^. **(q)** Extracellular TRAP activity of macrophages in osteoclastic induction culture supplemented with different concentrations of Mg^2+^. **(r)** The osteoclastic related gene expression of macrophage in osteoclastic induction culture supplemented with different concentrations of Mg^2+^.

**Fig. 6:**
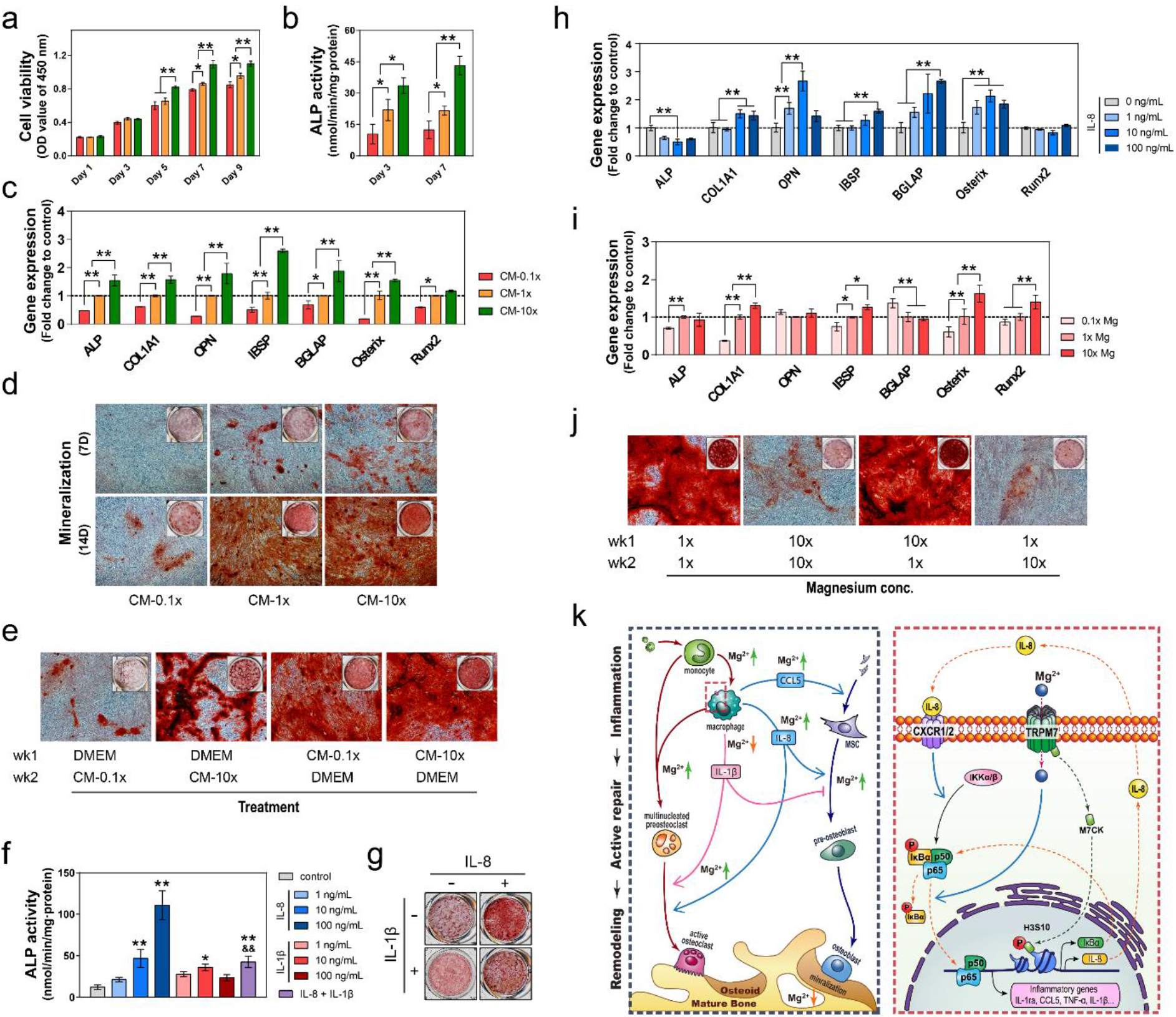
The effects of Mg and its modulated inflammatory microenvironment on osteogenesis. **(a, b, c)** The proliferation **(a)**, ALP activity **(b)** and osteogenic related gene expression **(c)** of MSC cultured in conditional medium from macrophages stimulated with different concentrations of Mg^2+^. *P*>0.05, \*\**P*<0.01 by one-way ANOVA with Tukey’s *post hoc* test. **(d)** Alizarin Red staining of mineralized nodules of MSC cultured in conditional medium from Mg-treated macrophages. **(e)** Alizarin Red staining of mineralized nodules of MSC treated with conditional medium from macrophages stimulated with 0.1x Mg or 10x Mg only at the first or second week of osteogenic induction. **(f, g)** The ALP activity **(f)** and the Alizarin Red stained mineralized nodules **(g)** of MSC cultured in medium supplemented with recombinant human IL-8 or IL-1β. **(h)** The osteogenic related gene expression of MSC cultured in different concentrations of recombinant human IL-8. **(i)** The osteogenic related gene expression of MSC cultured in DMEM supplemented with different concentrations of Mg^2+^. **(j)** Alizarin Red staining of mineralized nodules of MSC stimulated with 10x Mg at the different stages of osteogenic induction. **(k)** The schematic showing mechanism in which Mg^2+^ modulates both macrophages and mesenchymal stem cells in the bone healing processed.

To elucidate the role of TRPM7 in Mg^2+^-mediated inflammation modulation, we selectively silenced the expression of TRPM7 with siRNA (Fig. S4l) or blocked the channel activity with FTY720. Consequently, gene expression of inflammatory cytokines, such as IL-8, IL-1β, TNF-α and IL-1ra was dramatically increased to around ten-fold of the original level when the anti-inflammatory effect of Mg^2+^ was deprived (Fig. 5i, j). Moreover, as the nuclear translocation of M7CKs and the phosphorylation of Histone H3 were inhibited by the RNAi-knockdown or chemical blockage of TRPM7 (Fig. 5k), the effect of Mg^2+^ on regulating cytokine expressions, such as the induction of IL-8 and the downregulation of IL-1β, was completely abolished (Fig. 5i, j). In this study, we demonstrated that prolonged expose (i.e., 3 days or more) in Mg^2+^ also led to the increased expression of IKK-α and IKK-β, which contributed to the phosphorylation of IκB and p65, as well as the nuclear translocation of p65 (Fig. 4l, m). This suggests the stimulation of Mg^2+^ is associated with the activation of NF-κB signaling pathway in macrophages. Additionally, both siRNA knockdown or chemical blockage of TRPM7 suppressed the activation of NF-κB signaling pathway, by downregulating the expression of IκB kinases (IKK-α/β) and the phosphorylation levels of p65 and IκB in macrophages (Fig. 4n). Furthermore, our data showed that, with the supplement of RANKL and M-CSF, extended presence (i.e., 14 days or more) of Mg^2+^ resulted in an increased number of TRAP+ cells during the osteoclastic differentiation of THP1-derived macrophages (Fig. 5o, p), and increased extracellular TRAP activity (Fig. 5q). Interestingly, our RT-qPCR data revealed that only early osteoclastic markers (i.e., TRAP, RANK, and M-CSF) were elevated by the addition of Mg^2+^, whereas the late osteoclastic markers (i.e., CTSK and CTR), which indicates the maturation of osteoclasts were downregulated (Fig. 5r).

### Mg^2+^-mediates cytokines from macrophage contribute to bone formation

Since the macrophage-dependent inflammatory microenvironment was known to play an essential role in Mg^2+^-induced osteogenesis, we tested the hypothesis by treating human mesenchymal stem cells (MSC) with conditioned media (CM) harvested from THP1-derived macrophages pre-incubated in different concentrations of Mg^2+^ (named CM-0.1x, CM-1x and CM-10x). As expected, CM from 10x Mg^2+^-treated macrophages significantly promoted the proliferation (Fig. 6a) and osteogenic differentiation of MSC, as demonstrated by a two-fold increase in ALP activity (Fig. 6b) and the upregulation of osteogenic markers, including *ALP, COL1A1, OPN, IBSP, BGLAP*, and *Osterix* at both mRNA (Fig. 6c) and protein levels (Fig. S5f). Meanwhile, CM0.1x suppressed MSC proliferation and osteogenic gene expression (Fig. 6a-c). Alizarin red staining also revealed that CM from Mg^2+^-treated macrophage culture could accelerate the formation of mineralized nodules in MSC (Fig. 5g). Moreover, the osteogenic inducing effect of CM from Mg^2+^ stimulated macrophage was more pronounced when administered during the first week of osteogenic induction of MSC (Fig. 6e). Since our *in vitro* cytokine profiling (Fig. 4f-i) and *in vivo* immunostaining indicated IL-8 and IL-1β were two of the most significantly regulated cytokines from macrophages in response to the stimulation of Mg^2+^, we tested the effects of these two factors, individually and in combination, on MSC differentiation. The results indicated that human recombinant IL-8 replicated the effect of 10x Mg^2+^-stimulated CM, as evidenced by the significant increase in MSC proliferation (Fig. S5i), ALP activity (Fig. 6f), mineralization (Fig. 6g), and osteogenic genes expression (Fig. 6h). Although IL-1β led to a marginal increase in ALP activity and mineralization formation, it antagonized the osteogenic effects of IL-8 (Fig. 6f, g).

To distinguish the effect of macrophage-derived molecules from the direct action of Mg^2+^ on MSC, we also evaluated the osteogenic potential of MSC cultured in medium with different levels of Mg^2+^. Although Mg^2+^ insufficiency (0.1x) significantly reduced the ALP activity and the expression of osteogenic marker genes of MSC (Fig. 6i, S5b), an elevated level of Mg^2+^ (10x) only contributed to a <10% increase in the proliferation (Fig. S5a) and a marginal increase in early osteogenic gene markers, such as COL1A1, Osterix and Runx2 (Fig. 6i). Instead, most of the late osteogenic markers, such as OPN and BGLAP were not affected by the addition of Mg^2+^ (Fig. 6i). Moreover, even the Sirius red staining showed that 10x Mg^2+^ promoted the formation extracellular matrix (Fig. S5c), a prolonged (2 weeks) exposure of a high level of Mg^2+^ (10x) could suppress the spontaneous mineralization (Fig. 5j, S5d). Interestingly, by exposing MSC to high (10x) Mg^2+^ in different time windows, we noticed that the inhibitory effect of high Mg^2+^ on MSC mineralization was only observed when it was administered during the second half of the two-week culture experiment (Fig. 5j). To further explore the temporal effects of Mg^2+^ on osteogenic differentiation, we analyzed precipitation formed under simulated physiological condition using transmission electron microscopy (TEM). The typical TEM images and selected area electron diffraction (SAED) patterns of the precipitated calcium salts showed that the formation of crystallized apatite was significantly inhibited by the increase of Mg^2+^ level in the medium (Fig. S5g). X-ray diffraction data also indicated that the addition of Mg^2+^ led to a dramatic reduction of the hydroxyapatite phase, the predominant inorganic component in hard tissues (Fig. S5h).

## Discussion

Mg^2+^ has been extensively used for the modification of orthopedics biomaterials due to observations that Mg^2+^ can contribute to enhanced osteogenesis ^18,41^. Previous studies have speculated that Mg^2+^ modulates multiple signaling pathways at different stages in the differentiation process of mesenchymal stem cells toward osteoblast lineage ^24,41–43^. For example, Mg^2+^ is suggested to promote the proliferation and osteogenic differentiation of MSC by sequentially activating MAPK/ERK signaling pathway and Wnt/β-catenin signaling pathway ^43^. The synthesis of collagen type X (COL10A1), an extracellular matrix component involved in bone healing process, is reported to be enhanced by Mg^2+^ treatment through the nucleus translocation of hypoxia-inducible factors ^42^. Moreover, the activation of transforming growth factor-beta (TGF-β) and bone morphogenic protein (BMP) signaling, who plays fundamental roles in regulating MSC differentiation during skeletal development, is linked to the stimulation of Mg^2+^ ^23^. Although our *in vitro* data in this study verified the direct osteogenic action of Mg^2+^ on MSC, especially for the initiation of osteogenic differentiation, we noticed excessive Mg^2+^ didn’t contribute to late osteoblastic activities like the mineralization of extracellular matrix. Indeed, most of the studies suggest that the addition of Mg^2+^ needs to be tailored in an optimal range (around 2 mM – 8 mM) to achieve its osteogenic effects, otherwise, the presence of high levels of Mg^2+^ can be detrimental to the osteogenic differentiation of bone-forming cells ^18,42,44^. This means the conventional osteoblast lineage oriental study can not well address the osteogenic effect of Mg and its alloy, because the bone-forming cells are continuously challenged by high concentrations of Mg^2+^ in this scenario ^13,45^. Recently, implant-derived Mg^2+^ is demonstrated able to trigger neuronal calcitonin gene-related polypeptide-α release from dorsal root ganglia, resulting in enhanced osteogenic differentiation of periosteum-derived stem cells and improved bone-fracture healing ^46^. This finding suggests that Mg^2+^ may contribute to the osteogenic differentiation of osteoblast lineage through the activation of various other cell types in the bone microenvironment.

By grafting the Mg^2+^ releasing hydrogel at different stages of bone healing, our animal studies reveal there remains an effective window for the administration of Mg^2+^ to achieve better bone healing: the initial inflammation stage is more important than the subsequent active bone repair stage in term of the osteopromotive effect of Mg^2+^. Since macrophages are known for their dominant role in the innate inflammatory response and their regulatory effects on bone homeostasis in tissue regeneration ^38,47^, we asked whether the temporal effect of Mg^2+^ could be mediated through macrophage functions. By selective depletion of phagocytes in early stage of inflammation using liposome clodronate, we abolished the osteogenic effect of Mg^2+^. This suggests macrophages to be the primary target cells responsible for Mg^2+^-induced immunomodulation. Macrophages have been known to play important roles in the healing of many systems ^48^, including the healing of bone tissues. The depletion of macrophages is reported to contribute to complete loss of osteoblast-mediated bone formation at the modeling site, early skeletal growth retardation, and progressive osteoporosis ^49–51^. Activated macrophages were reported to locate at the sites of intramembranous bone deposition by forming an organized canopy structure over matrix-producing and -mineralizing osteoblasts ^51^.

In this study, our *in vivo* and *in vitro* data have provided compelling evidence showing the cellular activities of macrophages, especially the cytokine release profiles, to be significantly affected by the presence of Mg^2+^. First, the presence of Mg^2+^ facilitates the recruitment and activation of monocyte towards matured macrophage. Interestingly, although the upregulations of CD163 and CD206 suggest an M2 polarization of the Mg^2+^-treated macrophages, their cytokine secretion profile was distinct from the classical M1 and M2 regimes ^52^, as inflammatory genes typically observed in M1 macrophages (e.g., IL-1β and TNF-α) and in M2 macrophages (e.g., IL-10) were not significantly upregulated (Fig. 4i). Moreover, genes known to be involved in macrophage-mediated osteogenesis (e.g., *BMP2* ^*53*^, *OSM* ^*32*^, *TGF-β1* ^*54*^) were not dramatically altered by Mg^2+^ treatment. Instead, the Mg^2+^-triggered inflammatory activity demonstrated in our study, characterized by an increased level of CCL5, IL-8, and IL-1ra, might be distinct from other observations reported previously. CCL5 and IL-8 are traditionally recognized as pro-inflammatory factors responsible for cell recruitment to the sites of injuries and inflammation ^55,56^, however, both of them have been demonstrated capable of recruiting MSCs to bone healing site ^57,58^. Moreover, IL-8 has been shown as a crucial mediator for angiogenesis ^55,59^, the commitment of MSCs to bone regeneration ^57,60^, and restricting bone resorption activity of osteoclasts ^61^. IL-1β is one of the major pro-inflammatory cytokines that stimulate osteoclastogenesis ^62^. A reduction of IL-1β in response to the addition of Mg^2+^, possibly due to the anti-inflammatory effect of Mg^2+^ and the IKK-dependent activation of NF-κB ^63^, contributes to the regeneration of new bone. Additionally, it’s noteworthy that Mg^2+^ also upregulated the expression of IL-1 receptor antagonist (IL1-ra), which further limits IL-1β functions ^64^. Collectively, our observations suggest that Mg^2+^ stimulated macrophage to a cytokine mixture tailored for bone regeneration.

TRPM7 is a predominant Mg^2+^ channel in mammalian cells. Conditional gene deletion experiments suggest that this transporter is involved in a variety of immune-related functions, including macrophage activation and polarization in a current dependent manner ^65^, but the implications of these functions in tissue regeneration haven’t been well explored. Moreover, the intracellular Ser/ Thr kinase domain located at the carboxyl terminus of the membrane channel is reported able to modulate cellular behaviors via directly regulating the channel activity ^66,67^ or binding to multiple nuclear chromatin-remodeling complexes in nucleus ^68^. We found that the expression of TRPM7 in macrophages was intriguingly upregulated upon the stimulation of Mg^2+^, resulting in the increased intracellular level of Mg^2+^, as well as the cleavage and nuclear translocation of TRPM7 kinase fragments (M7CKs). Consequently, consistent with previous study ^68^, these chromatin-bound M7CKs directly contribute to the phosphorylation of Histone H3 at Ser 10 (p-H3S10), which has been reported to be critical for the recruitment of IKK to NF-κB signaling responsive promotors ^69^ and the binding of IKK to a variety of inflammatory gene promoters, such as CCL5 and IL-8 ^69–71^. Our ChIP assay demonstrated that Mg^2+^ led to an increased association of p-H3S10 with the promoters of IL-8 and IκBα, indicating the potent role of Mg^2+^ in activating NF-κB signaling and the production of inflammatory cytokines like IL-8.

The study demonstrates the essential role of TRPM7 in Mg^2+^-induced activation of NF-κB signaling pathway, which is crucial in inflammation responses ^72^. However, our experiments with the TRPM7 blocker FTY720 reveal that the channel activity of TRPM7 does not account for the entire spectrum of defects seen in TRPM7 siRNA treated macrophages, especially in terms of the phosphorylation of H3S10 and the activation of NF-κB signaling. This implies the cleaved kinase domain may play a more important role in the regulation of inflammatory cytokines in response to Mg^2+^. Recently evidence suggests that divalent cation like Ca2+ may play dual roles in macrophage through TRPM7: a rapid role in “jump starting” TLR4 endocytosis and possibly a slower role in tailoring an appropriate inflammatory response by modifying the activity of NF-κB ^73^. Therefore, we propose that the TRPM7-mediated inflammatory response to the stimulation of Mg^2+^ can be time-dependent: an initial M7CK oriented phosphorylation of H3S10 located at specific gene promoters followed by a better tailored activation of NF-κB signaling. This idea is supported by our observation that the promotive effect of Mg^2+^ on IL-8 secretion and the inhibitory effect on IL-1β was abrupted between 24 - 48 hours after the treatment. Moreover, since IL-8 also serves as a strong inducer for the activation of NF-κB signaling, the increased IL-8 in the microenvironment resulting from the M7CK-mediated histone modification can exaggerate the NF-κB signaling cascade through the IL-8 receptor (CXCR1/2) expressed on the membrane of macrophage ^74,75^. This explains why prolonged exposure in Mg^2+^ leads to unnecessary over activation of NF-κB signaling and osteoclastic differentiation of macrophages.

Unlike the TRAP+ multinucleated cells observed in bone remodeling stage, the TRAP+ multinucleated cells peaked on day 7 during bone healing are suggested to be responsible for the inflammatory response to biomaterials instead of bone resorption ^76^. These osteoclastic precursors present at early stage of bone healing are suggested to be actively involved in bone formation by the secretion of inflammatory cytokines and chemokines ^77^. We noticed that a short-term delivery of Mg^2+^ contributed to an increased number of TRAP+ preosteoclasts at inflammation stage but resulted in a decreased number of bone-resorbing osteoclasts in remodeling stage. By contrast, when the delivery of Mg^2+^ was extended, the protective effect of Mg^2+^ from osteoclastogenesis at remodeling stage disappeared. Given the above-mentioned influence of Mg^2+^ on the activation of NF-κB signaling, which plays a central role in RANK ligand-induced osteoclastogenesis ^78^, the administration of Mg^2+^ besides early inflammation stage may be detrimental for bone regeneration.

The formation of calcified tissue in the biological environment starts with the nucleation and growth of calcium phosphate precursors, however, the presence of Mg^2+^ may inhibit this process by selectively stabilizing more acidic hydrated precursor phases, like dicalcium phosphate ^79,80^. Moreover, the adsorption of excessive Mg^2+^ on the surface of precursor and nuclei could further lead to the blocking of the active growth sites 81. During the investigation of the direct effect of Mg^2+^ on MSC, we observed that a high level of Mg^2+^ (10x) at the beginning of the osteogenic induction of MSC towards matured osteoblast could promote extracellular matrix formation and mineralization. However, when 10x Mg^2+^ was supplemented at late stage or throughout the osteogenic induction, the mineralization of MSC could be dramatically impaired. This finding recapitulated our *in vivo* data showing that prolonged Mg^2+^ treatment could reduce BMD, as extensively reported elsewhere ^12,46^. We interpret these data as an indication of the biphasic nature of Mg^2+^ effect on biological mineralization: Mg^2+^ and the osteogenic factors secreted by macrophage upon the stimulation of Mg^2+^ are effective during the early phase of osteogenic differentiation of MSC. However, the MSC committed to osteoblast lineage becomes less responsive to Mg^2+^ and the Mg^2+^-induced immune factors. Instead, the persistent presence of Mg^2+^ beyond this stage suppresses mineralization.

## Conclusion

This study establishes a new paradigm for understanding the diverse and multifaceted roles of Mg^2+^ in bone healing (summarized in Fig. 6k). The delivery of Mg^2+^ contributes to the infiltration and the activation of the monocyte-macrophage-preosteoclast lineage in the bone defect. The influx of Mg^2+^ through TRPM7 channel and the nuclear translocation of M7CKs lead to the polarization of the macrophages into a pro-osteogenesis subtype facilitating the recruitment and the osteogenic differentiation of MSC, which is distinct from the classical M1 or M2 phenotypes. Our findings suggest that the functional plasticity of macrophages may be even more prevalent than previously described, eliciting specific responses tailored for different tissue or injury types. It is possible that the initial immune response to bone injury, orchestrated by cell types in the monocyte-macrophage-preosteoclast lineage, can be tuned to support tissue regeneration by a variety of signals, with Mg^2+^ being a defining example. However, the osteo-promoting functions of Mg^2+^ only operate during the early phase of osteogenesis as continued stimulation of Mg^2+^ leads to over activation of NF-κB signaling and the osteoclastic differentiation of macrophage. Moreover, the presence of Mg^2+^ in active bone repair/remodeling stage inhibits the calcification of extracellular matrix, resulting in compromised bone formation. In light of our findings, the use of Mg^2+^-based biomaterials as orthopedic implants must be viewed with caution, as these materials usually sustain an extended release of Mg^2+^ which permeates into the wound with uncontrolled dose kinetics. The delicate balance between the osteo-promoting and osteo-inhibitory effects of Mg^2+^, if unchecked, will lead to unpredictable clinical outcomes.

## Methods

### Preparation of alginate gel

20% alginate gel (Alg) was prepared by mixing sodium alginate powder (Sigma-Aldrich, USA) in deionized water, while 10% magnesium chloride (MgCl_2_, Sigma-Aldrich) was used for the preparation of magnesium cross-linked alginate (Mg-alg). The gel was then loaded in syringe and sterilized by gamma rays (Co-60) irradiation at a dose of 25kGy. The as-prepared gel was immersed in 1X Phosphate-buffered saline (PBS) at a ratio of 0.1 g/mL and kept in an incubator at 37 ℃ for five days. At designated time point, the Mg^2+^ concentration in the PBS was measured by inductively coupled plasma optical emission spectrometry (ICP-OES, Optima 2100DV, Perkin Elmer, USA).

### Animal surgery

All the animal procedures were performed in accordance with a protocol approved by the Committee on the Use of Live Animals in Teaching and Research (CULATR, HKU). 6-8-week old Sprague Dawley (SD) rats, weighting 200-250 g, were purchased from Charles River Lab (USA) and maintained in specific pathogen-free facility (Lab animal unit, LAU, HKU). After the rats were anesthetized via intraperitoneal injections of ketamine hydrochloride (67 mg/kg; Alfamine, Alfasan International B.V., Holland) and xylazine hydrochloride (6 mg/kg; Alfazyne, Alfasan International B.V.). A tunnel defect, which is 2 mm in diameter, was prepared at the lateral epicondyle of each rat using a hand driller. Subsequently, the defect was carefully filled with either pure alginate or Mg-crosslinked alginate. After the layer by layer closure of wound, the rats immediately received subcutaneous injections of terramycin (1 mg/kg) and ketoprofen (0.5 mg/kg).

### Micro-CT analysis

In order to monitor the healing process and examine new bone formation around the gel, micro-CT scans were conducted at serial time points after the surgery (*i.e.*, 0, 7, 14, 21, 28 and 56 days). After the rats were anesthetized, the grafted site was scanned using a live animal scanning device (SkyScan 1076, Kontich, Belgium). Two phantom contained rods with standard density of 0.25 and 0.75 g/cm^3^ were scanned with each sample for calibration. The resolution was set at 17.3 μm/pixel. Data reconstruction was done using the NRecon software (Skyscan Company), and image processing and analysis were done using CTAn software (Skyscan Company).

### Fluorochrome labeling

Two fluorochrome labels were used sequentially to evaluate bone regeneration and remodeling within the defects. Calcein green (5 mg/kg, Sigma-Aldrich) was subcutaneously injected into rat femora one week after the surgery, while xylenol orange (90 mg/kg, Sigma-Aldrich) was injected three weeks after the surgery. The fluorochrome labels were visualized under a fluorescence microscopy (Niko ECL IPSE 80i, Japan). The intensity of fluorescence was analyzed by ImageJ software (NIH, USA).

### Histological analysis

After the execution of the rats, the femora were collected and fixed in 4% paraformaldehyde solution. For non-decalcified sections, the samples were dehydrated with gradient ethanol and embedded with methyl-methacrylate (MMA, Technovit 9100 New, Heraeus Kulzer, Hanau, Germany) after using xylene as a transition. Then, the embedded samples were cut into slices with a thickness of 200 μm and micro-ground to a thickness of 50-70 μm. The selected sections were stained with Goldner’s trichrome (Sigma-Aldrich) staining. For decalcified sections, the samples were decalcified with 12.5% ethylenediaminetetraacetic acid (EDTA, Sigma-Aldrich) for 6 weeks. The specimens were then dehydrated in ethanol, embedded in paraffin, and cut into 5 μm-thick sections using a rotary microtome (RM215, Leica Microsystems, Germany). Haematoxylin and eosin (H&E) staining and TRAP staining (sigma-Aldrich) was performed in selected slides from each sample following manufacturer’s instructions. Images were captured using a polarizing microscopy (Nikon Eclipse VL100POL, Nikon, Tokyo, Japan)

### Scanning electron microscopy-energy dispersive spectroscopy (SEM-EDS)

MMA embedded bone specimens were sliced and polished to ~100 μm before observation. The elemental composition and distribution in the grafted area were determined by SEM-EDS (SU1510, Hitachi, Tokyo, Japan).

### Immunohistochemistry analysis

The dewaxed slide was treated with Proteinase K (Sigma-Aldrich) for proteolytic digestion and 3% H_2_O_2_ for the elimination of endogenous peroxidase activity. After blocking with normal goat serum, the slides were incubated with primary antibody overnight at 4°C, respectively. The primary antibodies used in this study include: rabbit anti-OCN (Abcam), rabbit anti-IL-8 (Abcam), rabbit anti-CCL5 (Abcam), rabbit anti-IL-1β (Abcam), rabbit anti-IL-1ra (Abcam), anti-CD68 (Abcam) and rabbit anti-TRPM7 (Abcam). The slides were then incubated with goat anti-rabbit secondary antibody and visualized using Diaminobenzidine (DAB) staining kit (Santa Cruz Biotechnology, Santa Cruz, USA) following the manufacturer’s instruction. Immunofluorescent staining was done using Alexa-Fluor 488 conjugated secondary antibody (Thermo Fisher Scientific) and Hoechst 33324 (Thermo Fisher Scientific).

### Detection of inflammatory cytokines

The femora of the rats grafted with pure alginate or Mg-crosslinked alginate were harvested. The bone tissues in and around the grafted defects, around 1 cm in length, was ground into mud using a ceramic mortar and pestle. The mud of bone tissue was then homogenized in pre-cooled RIPA Lysis and Extraction Buffer (ThermoFisher Scientific) for 1 h. The buffer solution was centrifuged at 15,000 rpm for 20 min at 4°C. The supernatant was collected for protein concentration quantification with BCA Protein Assay Kit (ThermoFisher Scientific). An equal amount of protein from each sample was subjected to quantitative analysis of IL-8 and IL-1β using ELISA kit (R&D system) following the manufacturer’s instructions.

### Cell culture

The human monocyte cell line THP1 was obtained from the American Type Culture Collection (ATCC, VA, USA), and maintained in RPMI 1640 supplemented with 10% heat-inactivated fetal bovine serum (FBS, Thermo Fisher Scientific) and 1% (v/v) penicillin/streptomycin (Thermo Fisher Scientific) at 37℃ under 5% humidified CO_2_. Human mesenchymal stem cells (MSC) was kindly provided by Prof. D. Campana (St Jude Children’s Research Hospital, Memphis, Tennessee), and maintained in Dulbecco’s modified Eagle Medium (DMEM, Gibco) supplemented with 10% FBS (Thermo Fisher Scientific) and 1% (v/v) penicillin/streptomycin (Thermo Fisher Scientific) at 37 ℃ under 5% humidified CO_2_.

Macrophages were differentiated from THP1 cells using serum-free DMEM supplemented with 10 ng/mL phorbol 12-myristate 13-acetate (PMA, Sigma-Aldrich). After 48h, the THP1-derived macrophages were further cultured in DMEM containing different concentrations of magnesium (i.e., 0.08 mM, 0.8 mM, and 8 mM) for another 72 h to allow full differentiation and polarization. For osteogenic differentiation of MSC, osteogenic supplements (5 mM β-glycerol phosphate, 0.05 mM L-ascorbic acid 2-phosphate and 10^−7^ M Dexamethasone) were added to the serum-free DMEM.

### Flow Cytometry assay

Flow cytometric assay was performed using FACSCantoII Analyzer (BD Biosciences, USA). Differentiated macrophages were detached with trypsin and washed with 1X PBS. For the detection of macrophage surface markers, cells were incubated with monoclonal mouse anti-human antibodies CD163-Cy5.5, CD206-FITC, and CD80-FITC (BD Biosciences, USA), or relevant isotypes (BD Biosciences) for 1 hr at 4 ℃ in dark. After washing with 1X PBS, the fluorescence was compared to isotypes with 10,0000 events recorded. All flow cytometric data was analyzed using Flowjo software, version 10 (Tree Star, USA).

### Cell attachment and proliferation assay

The effect of Mg^2+^ on the attachment of THP1-derived macrophages and the proliferation of MSC was assessed using a cell counting kit-8 (CCK-8, Dojindo, Japan). Cell viability at the designated time points was presented by the optical density (OD) value measured at the wavelength of 450 nm using a microplate spectrophotometer (SpectraMax 340, Molecular Devices, USA).

### ATP assay

After the stimulation of Mg^2+^, the macrophages were lysed, and the intracellular ATP concentrations were determined using a luminescence ATP detection assay (ATPLite, PerkinElmer, USA) following the manufacturer’s instructions.

### Cytokine assay

The cytokines produced by THP1-derived macrophages after the stimulation of different concentrations of Mg^2+^ were determined by Proteome Profiler antibody arrays (R&D System, USA) following the manufacturer’s instructions. The concentration of IL-8 and IL-1β was further confirmed by specific ELISA kits (R&D System).

### Alkaline phosphatase (ALP) assay

The effects of Mg^2+^ or conditional medium from macrophages on the ALP activity of MSC were evaluated using the p-NPP method. At the designated time points, the cells were lysed with 0.2% Triton X-100 at 4°C for 2 h. The supernatant of the lysis after centrifugation was collected and assayed using ALP detection kit (Sigma-Aldrich) following the manufacturer’s instructions. The total protein content was measured using BCA Protein Assay Kit (ThermoFisher Scientific). The relative ALP activity was normalized to total protein content and expressed as units/g·protein.

### Mineralization assay

Sirius Red staining was used to study the formation of extracellular matrix of MSC. At the designated time points, cells were fixed with 75% ethanol, and the collagen was stained with Sirius Red solution (Sigma-Aldrich) for 1 hour. Alizarin Red staining was used to study the mineralization of MSC. At the designated time points, cells were fixed with 4% paraformaldehyde, and the mineralization nodules were stained with Alizarin Red solution (Sigma-Aldrich) for 5 min. After a thorough wash with Millipore water, the sample was dried in air before photo taking.

### Real-time quantitative PCR (RT-qPCR) assay

The total RNA of the cells was extracted and purified using RNeasy Plus kit (Qiagen, USA) following the manufacturer’s instructions. For the reverse transcript, complementary DNA was synthesized using Takara RT master Mix (Takara, Japan) following the manufacturer’s instructions. The primers used in the RT-qPCR assay were synthesized by Life Technologies (ThemoFisher Scientific) based on sequences retrieved from Primer Bank (http://pga.mgh.harvard.edu/primerbank/, Table S1). SYBR Green Premix Ex Taq (Takara) was used for the amplification and detection of cDNA targets on a StepOne Plus Real-time PCR system (Applied Biosystems, USA). The mean cycle threshold (Ct) value of each target gene was normalized to the housekeeping gene GAPDH. The results were shown in a fold change using the ΔΔCt method.

### Western blotting

At the designated time points after the treatment, the cells were rinsed with ice-cold PBS and lysed with RIPA Lysis and Extraction Buffer (ThermoFisher Scientific). After centrifugation at 15,000×g for 10 min at 4°C, the supernatants were collected for measuring the protein concentration with BCA Protein Assay Kit (ThermoFisher Scientific). Cytosolic and nuclear extracts were prepared using NE-PER reagents (ThermoFisher Scientific) following the manufacturer’s instructions. A total of 30 μg of protein from each sample were subjected to SDS-PAGE electrophoresis and transferred to PVDF membrane (Merck Millipore, USA). Then the membrane was blocked in 5% w/v bovine serum albumin (BSA, Sigma-Aldrich) and incubated with blocking buffer diluted primary antibodies overnight at 4°C. The primary antibodies used include rabbit anti-TRPM7 (Abcam), rabbit anti-IL-8 (Abcam), rabbit anti-CCL5 (Abcam), rabbit anti-IL-1β (Abcam), rabbit anti-IL-1ra (Abcam), rabbit anti-OPN (Abcam), mouse anti-ALP (Santa Cruz, USA), rabbit anti-OCN (Abcam), rabbit anti-IKKβ (CST), mouse anti-IKKα (CST), rabbit anti-NF-κB p65 (CST), rabbit anti-Phospho-IκBα (CST), mouse anti-IκBα (CST), rabbit anti-Phospho-Histone H3S10 (Abcam), rabbit anti-Histone H3 (Abcam), and mouse anti-β-actin (CST, USA). The protein bands were visualized by ECL substrate (ThermoFisher Scientific) and exposed under ChemiDoc XRS System (BioRad, USA).

### Immunocytochemistry analysis

Following the stimulation of DMEM containing different concentrations of Mg^2+^, cells were washed with 1X PBS three times, permeabilized with 0.2% Triton X-100 for 10 min either before or after fixation with 4% paraformaldehyde. Then, the cells were incubated with rabbit anti-TRPM7 or rabbit anti-NF-κB primary antibody overnight, and stained with Alexa-Fluor 555 conjugated secondary antibody (Thermo Fisher Scientific), FITC-Phallotoxins (Sigma-Aldrich) and Hoechst 33342 (ThermoFisher Scientific) before observation using a Carl Zeiss LSM 780 confocal microscopy (Carl Zeiss, Germany).

### Detection of Mg uptake in macrophages

THP1-derived macrophages were incubated in Mg^2+^-free DMEM (ThermoFisher Scientific) supplemented with 4 μM Mag-Fluo-4 (ThermoFisher Scientific) and 1 μM F-127 (ThermoFisher Scientific) for 30 min. After that, cells were washed with Mg^2+^-free DMEM and counterstained with Hoechst 33324. Real-time images were captured using a spinning confocal microscope (Perkin Elmer Instruments, USA) immediately after the addition of 8.0 mM MgCl2 into the Mg^2+^-free DMEM. For visualization of mitochondrial, cells were pretreated with MitoTracer (ThermoFisher Scientific) for 1.5 hrs before observation.

### Super-resolution microscopy

THP1-derived macrophages were seeded on coverslips with locking beads. After the treatment, cells were washed with 1X PBS three times, fixed with 4% paraformaldehyde, and permeabilized with 0.2% Triton X-100. Then, the cells were incubated with mouse anti-TRPM7 and rabbit anti-Histone H3 primary antibodies overnight. After thorough washing steps with PBS, they were stained with Alexa-Fluor 647 and Alexa-Fluor 488 conjugated secondary antibody (Thermo Fisher Scientific) before being mounted to a Stochastic Optical Reconstruction Microscope (STORM, Nanobioimaging Ltd., Hong Kong).

### TRPM7 inhibition and blockage

For TRPM7 silencing, THP1-derived macrophages were transfected with 10 nM siRNA targeting human TRPM7 (SR310261, OriGene, USA) following the manufacturer’s instructions using siTran1.0 (OriGene, USA) as the agent. Cells transfected with nonspecific control siRNA (SR30004, OriGene, USA) were used as the control. siRNA transfection efficiency was verified 72 h after the transfection by western blots. For the inhibition of TRPM7 activity, cells were pretreated with 3 μM FTY720 (Sigma-Aldrich) for 2 h, washed with 1X PBS, and subjected to the other assays.

### Chromatin immunoprecipitation

After the stimulation, chromatin immunoprecipitation (ChIP) assay was performed using a ChIP kit (Abcam) as described by the manufacturer’s instruction. In brief, chromatin from crosslinked macrophages was sheared by sonication (8 of 30s-on and 30s-off pulses at high power output, Bioruptor Plus, USA) and incubated overnight with rabbit anti-Phospho-Histone H3S10 (Abcam) at 4°C. Precipitated DNAs were analyzed by RT-qPCR with SYBR Green Premix Ex Taq (Takara) and primers for human IL-8 promoter (−121 to +61) 5’-GGGCCATCAGTTGCAAATC-3’ and 5’-TTCCTTCCGGTGGTTTCTTC-3’, human IκBα promoters (−316 to −15) 5’-GACGACCCCAATTCAAATCG-3’ and 5’-TCAGGCTCGGGGAATTTCC-3’, as well as human β-actin promoter (−980 to −915) 5’-TGCACTGTGCGGCGAAGC-3’ and 5’-TCGAGCCATAAAAGGCAA-3’.

### Statistical Analysis

Each experiment was performed at least three times, and the results were expressed as means ± standard deviations (SD). For the data analysis, either one-way analysis of variance (ANOVA) followed by Tukey’s multiple-comparison post hoc test or two-sample *t*-test was performed with SPSS ver.13.0 (IBM SPSS, USA). The level of significant difference among groups was defined and noted as * p < 0.05 and ** p<0.01.

## Acknowledgments

We acknowledge HKU Li Ka Shing Faculty of Medicine Faculty Core Facility for providing a harmonious working environment. This work was financially supported by the General Research Fund of Hong Kong Research Grant Council (#17214516, #N_HKU725/16), Hong Kong Innovation Technology Fund (#ITS/147/15, ITS/287/17), Hong Kong Health and Medical Research Fund (#03142446), HKU Seed Fund for Translational and Applied Research (#201611160006), Sanming Project of Medicine in Shenzhen “Team of Excellence in Spinal Deformities and Spinal Degeneration Disease” (SZSM201612055), National Natural Science Foundation of China (No. 31370957, No.81470783), Shenzhen Science and Technology Funding (JCYJ20160429190821781& JCYJ20160429185449249) and Guangdong Scientific Plan (2014A030313743). We thank Dr. Stuart Fraser, School of Medical Sciences, University of Sydney, Australia and Prof. Cao Xu, Department of Orthopedics, School of Medcine, the Johns Hopkins University, for their useful comments.

## Author contributions

W. Qiao, K.H.M. Wong, J. Shen and W. Wang conducted animal surgery and analyzed the results. W. Qiao, J. Wu and J. Li contributed to the cell culture and the *in vitro* tests. K.H.M. Wong and Z. Lin were responsible for the preparation of Mg-releasing alginate. Z.T. Chen, K. Lai and Y.W. Lam contributed to the design of *in vitro* experiments. J.P. Matinlinna, Y. Zheng, S. Wu and X. Liu provide insightful comments on the material-science related issues. K.M.C. Cheung and Z.F. Chen contributed to the design of animal models and provided invaluable suggestions about clinical indications of the study. Y.W. Lam, K.W.K. Yeung and Z.F. Chen contributed to data interpretation and supervised the project. W. Qiao, Y.W. Lam and K.W.K. Yeung wrote the manuscript with input from all authors.

---

**Fig. S1:**
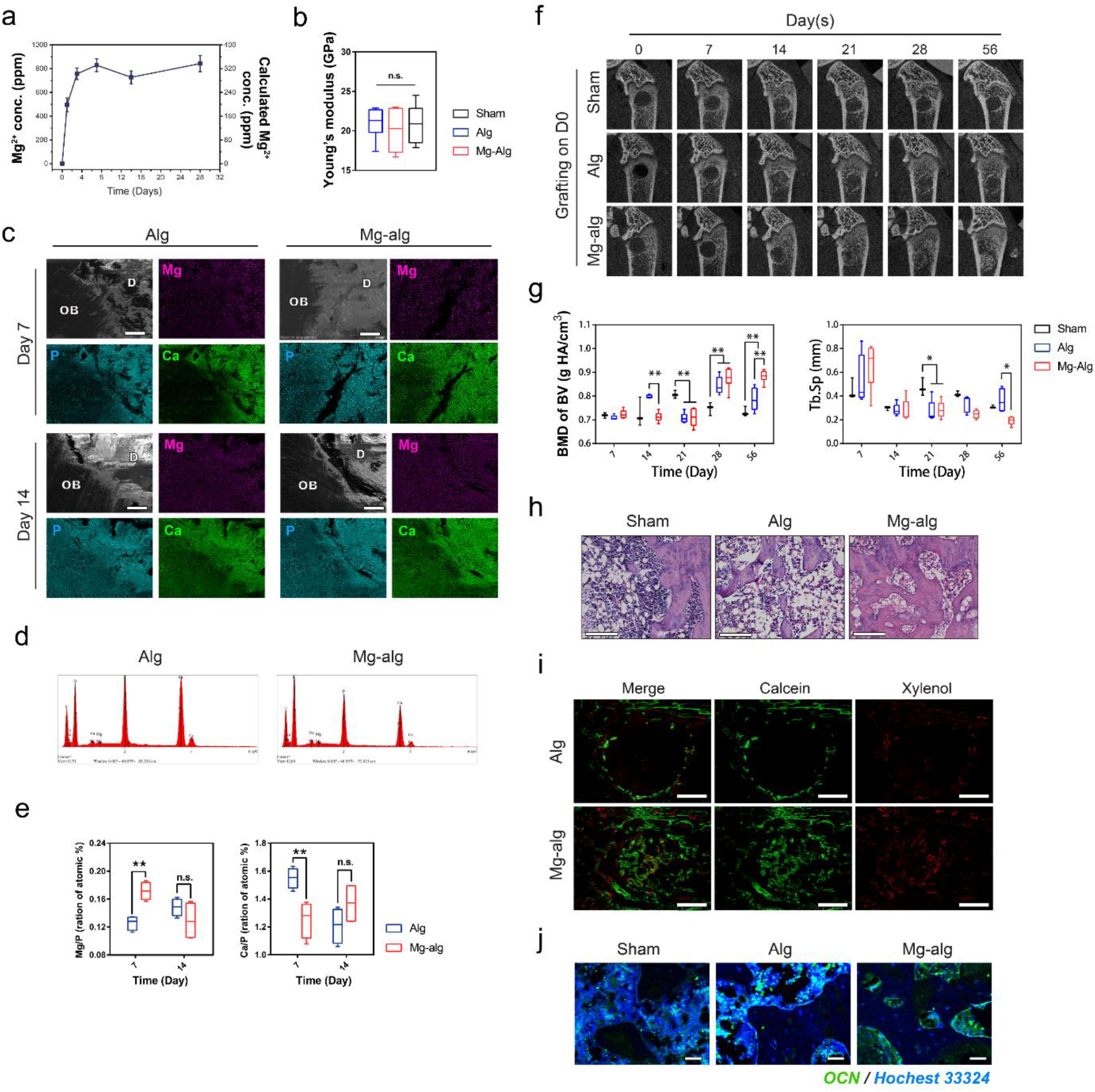
**(a)** Cumulative Mg^2+^ release from the Mg crosslinked alginate measured *in vitro* using ICP-OES. **(b)** Young’s modulus of newly formed bone in the sham group (n = 5), alginate group (n = 5) and Mg-alginate group (n = 5). **(c)** Representative SEM images and EDX mapping of Mg, P and Ca in and around the rat femoral defects grafted with alginate or Mg-releasing alginate 7 and 14 days after the operation, scale bars = 200 μm. **(d)** Representative SEM-EDX spectrums showing the atomic composition in the femur defects grafted with alginate or Mg-releasing alginate 7 days after the operation. **(e)** Quantitative data showing the Mg/P and CaP ratio in defect areas grafted with alginate or Mg-releasing alginate on day 7 and day 14. **(f, g)** Representative micro-CT images **(f)** and Corresponding measurements of BMD of BV and Tb.Sp. **(g)** showing the healing process of rat femoral defects without grafting (Sham group), grafted with pure alginate (serve as a control) or Mg-releasing alginate. **(h)** Representative H&E staining of the defects in rat femur on day 56, scale bars =200 μm. **(i)** Representative images of calcein/xylenol labeling for bone regeneration in the rat femoral defects grafted with alginate or Mg crosslinked alginate, scale bars = 1 mm. **(j)** Representative immunofluorescent images showing the expression of OCN in the grafted defects in the rat femora, scale bars =100 μm.

**Fig. S2:**
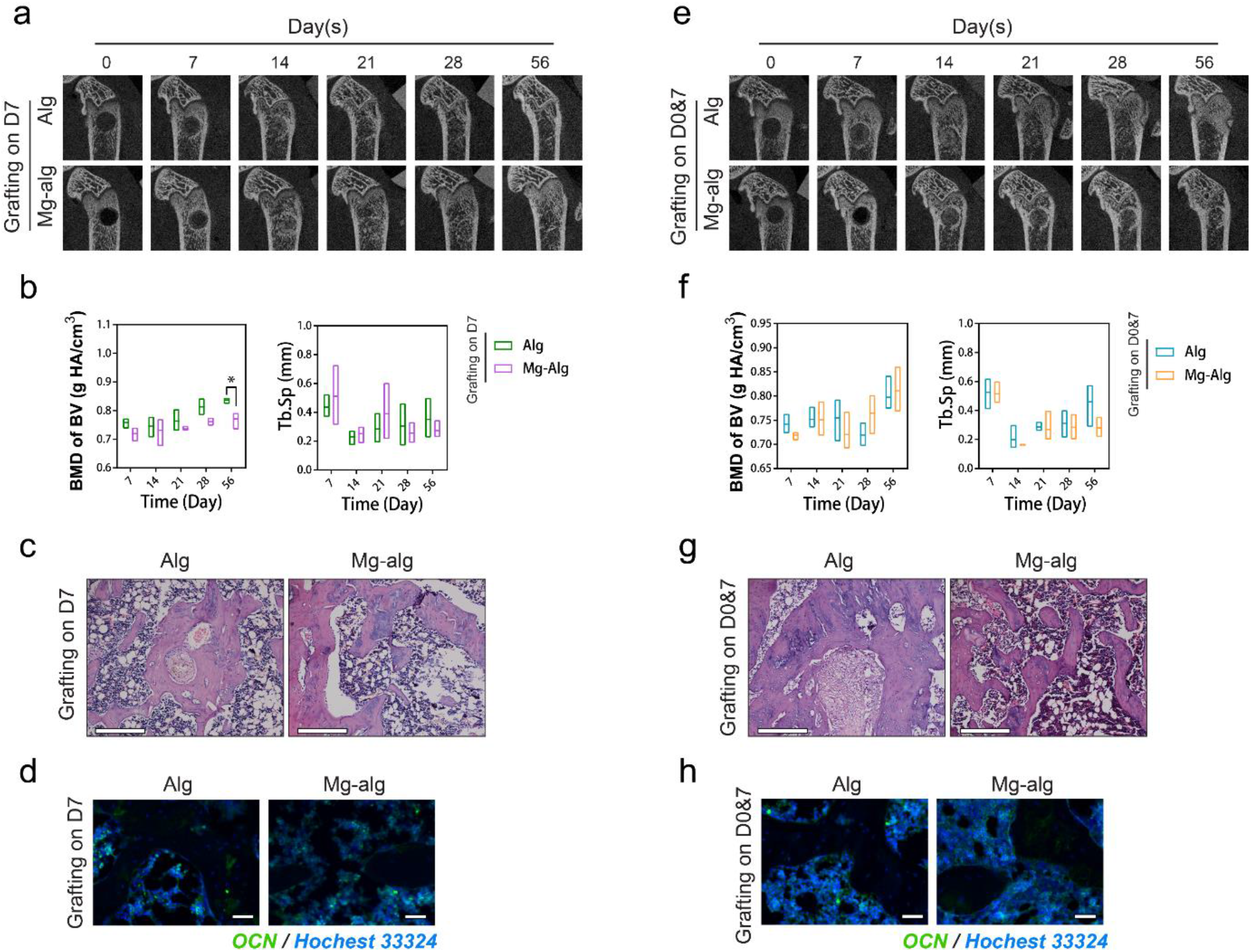
**(a, b)** Representative micro-CT images **(a)** and Corresponding measurements of BMD of BV and Tb.Sp **(b)** showing the healing process of rat femoral defects when pure alginate (serve as a control) or Mg crosslinked alginate were grafted on Day 7 after the operation. Data are mean ± s.d. \**P*<0.05, \*\**P*<0.01 by one-way ANOVA with Tukey’s *post hoc* test. **(c,d)** Representative H&E staining **(c,** scale bars =200 μm) and immunofluorescent images showing the expression of OCN **(d,** scale bars =100 μm) in rat femur defect when the material was grafted on Day 7 after the operation. **(e, f)** Representative micro-CT images **(e)** and Corresponding measurements of BMD of BV and Tb.Sp. **(f)** showing the healing process of rat femoral defects when pure alginate (serve as a control) or Mg crosslinked alginate were grafted both immediately and 7 days after the operation. **(g, h)** Representative H&E staining **(g,** scale bars =200 μm) and immunofluorescent images showing the expression of OCN **(h,** scale bars =100 μm) in rat femur defect when the material was grafted on Day 7 after the operation.

**Fig. S3:**
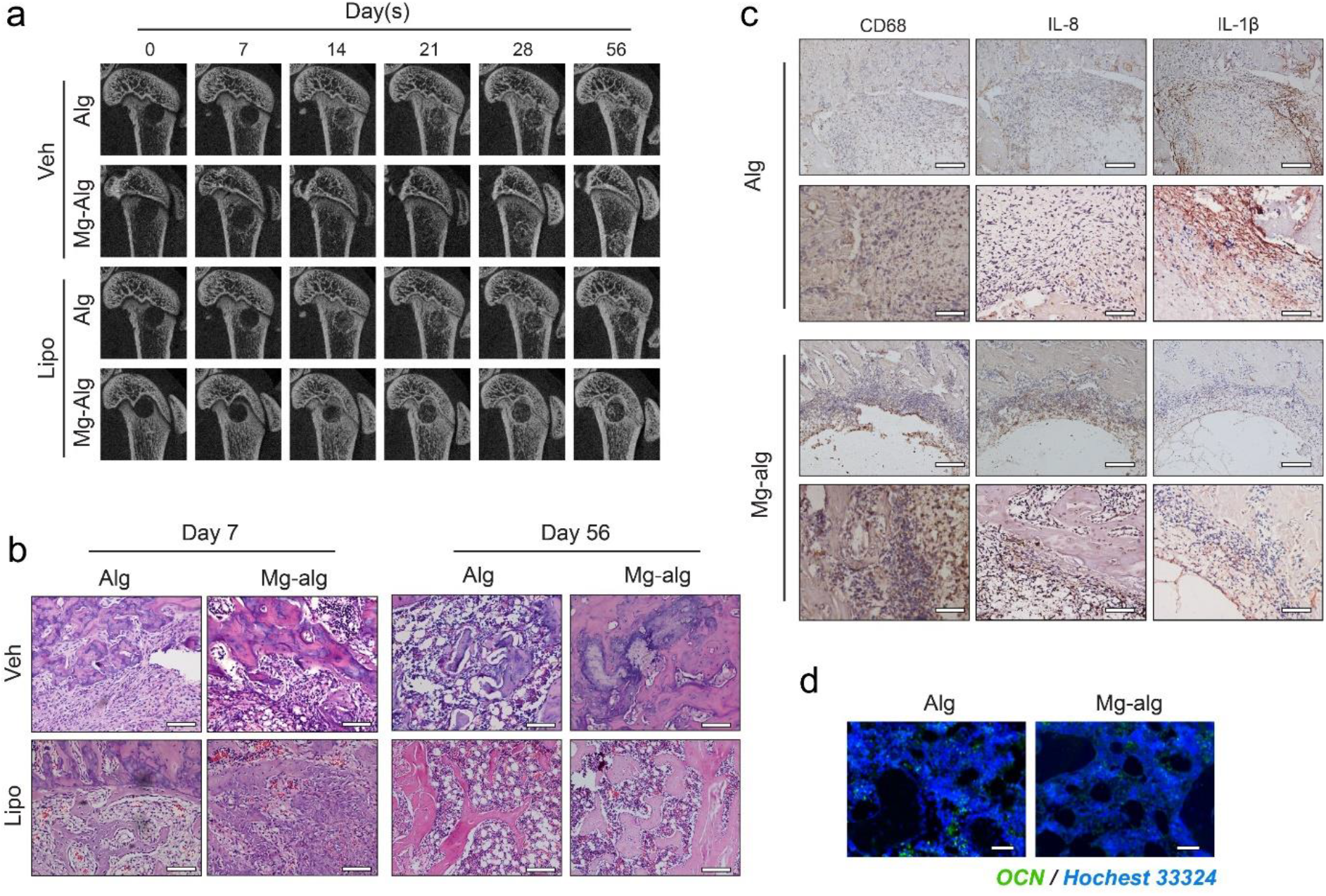
**(a)** Representative micro-CT images showing the healing process of femoral defects grafted with pure alginate (n = 3) or Mg-releasing alginate (n = 3) in macrophage depleted rats compared with control group injected with vehicle from day 7 to day 56. **(b)** Representative H&E staining of the defects in rat femur on day 7 and day 56 in control rats and macrophage depleted rats, scale bars =200 μm. **(c)** Representative IHC images showing the expression of CD68, IL-8 and IL-1β within the rat femoral defects grafted with alginate or Mg-alginate on day 7. Lower images (scale bar = 100 μm) are high-resolution versions of the upper images (scale bar = 200 μm). **(d)** Representative immunofluorescent images showing the expression of OCN in the grafted defect of macrophage depleted rats, scale bars =100 μm.

**Fig. S4:**
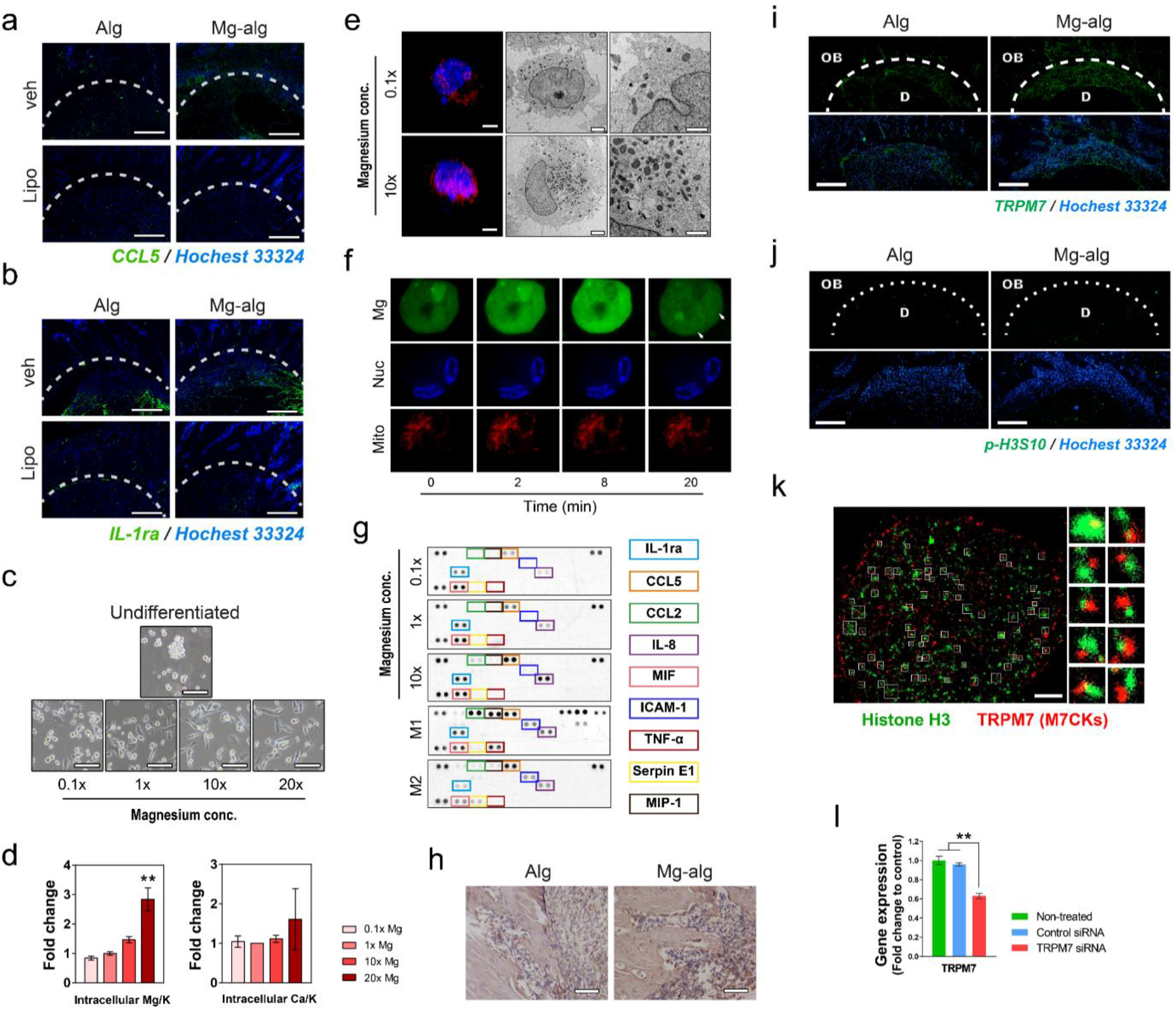
**(a,b)** Representative immunofluorescent images showing the expression of CCL5 **(a)** and IL-1ra **(b)** on day 7 in the grafted defects in the rat femora, scale bars =500 μm. **(c)** Representative microscopy images showing changes in the morphology of THP-1 derived macrophages after PMA-induced differentiation with the addition of different concentrations of Mg^2+^. **(d)** ICP-OES measurement of intracellular Mg to K ratio and Ca to K ration in macrophages after the stimulation of different concentrations of Mg^2+^. **(e)** Representative 3D confocal (scale bar = 2 μm) and TEM images showing the increased number of mitochondria in THP1-derived macrophages after the stimulation of Mg^2+^. right images (scale bar = 500 nm) are high-resolution versions of the middle images (scale bar = 2 μm). **(f)** Representative fluorescence images showing the entry of Mg^2+^ into macrophages, the accumulation of Mg^2+^ in nuclei, as well as the dynamic changes in mitochondria after the addition of 8 mM MgCl2. **(g)** Cytokine array showing the major cytokines produced by Mg^2+^ treated macrophages, as well as classical differentiated M1 and M2 macrophages. **(h)** Representative IHC images showing the expression of TRPM7 in the rat femoral defects grafted with alginate or Mg-alginate (scale bar = 100 μm). **(i, j)** Representative immunofluorescent images showing the expression of TRPM7 **(i)** and the phosphorylation of Histone H3S10 **(j)** in and around the defects grafted with alginate or Mg-releasing alginate 7 days after the surgery, scale bars = 200 μm. **(k)** Representative super-resolution images showing the colocalization of TRPM7 (M7CKs) and Histone H3 within the nucleus of the macrophage. **(l)** The effects of TRPM7 siRNA on the expression of TRPM7 in THP1-derived macrophages.

**Fig. S5:**
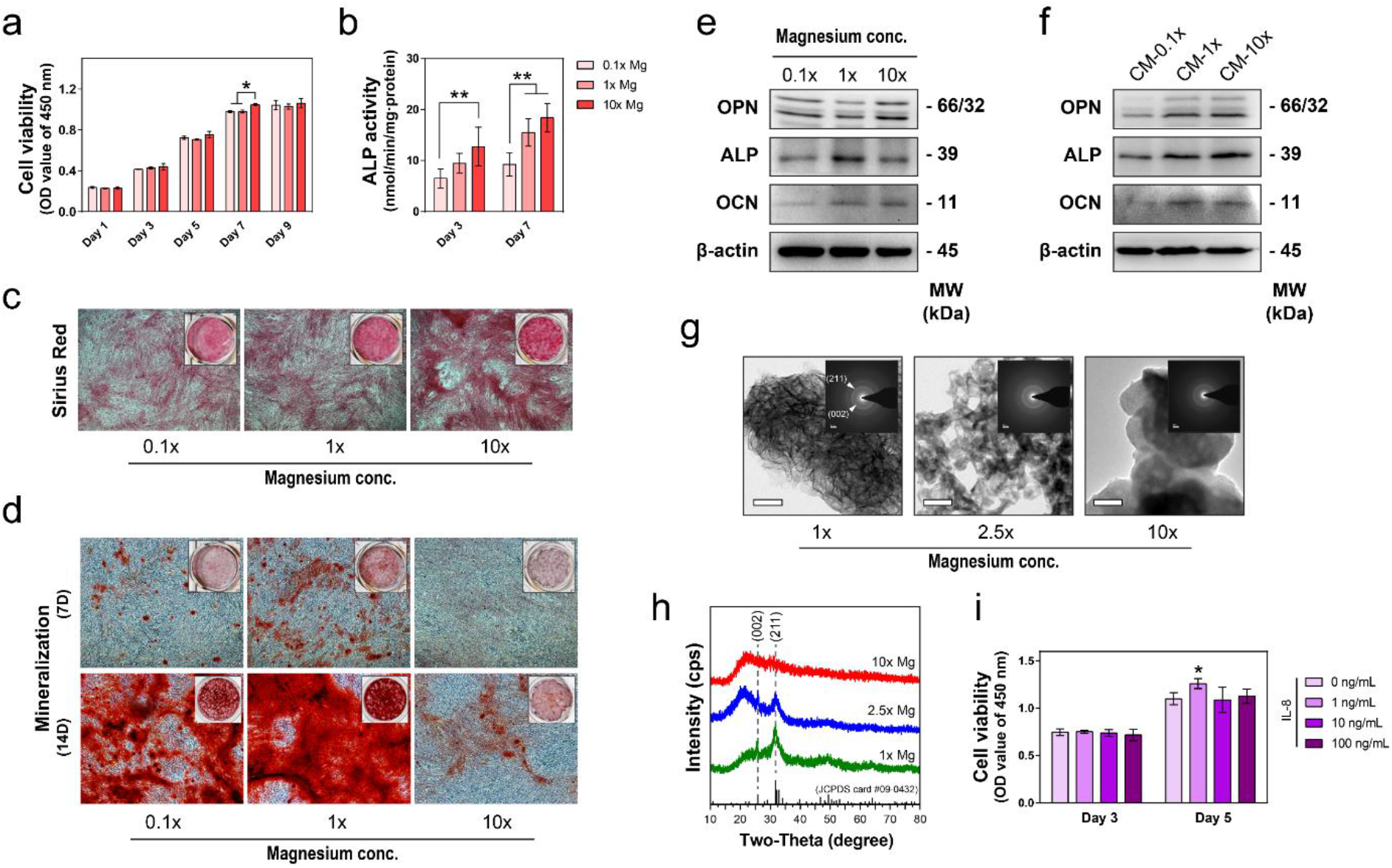
**(a, b)** The proliferation **(a)** and ALP activity **(b)** of MSC cultured in DMEM supplemented with different concentrations of Mg^2+^. Data are mean ± s.d. \**P*<0.05, \*\**P*<0.01 by one-way ANOVA with Tukey’s *post hoc* test. **(c)** Sirius Red staining showing collagen formation of MSC cultured in DMEM supplemented with different concentrations of Mg^2+^. **(d)** Alizarin Red staining of mineralized nodules of MSC cultured in DMEM supplemented with different concentrations of Mg^2+^. **(e, f)** Representative western blots showing the bone markers expression of MSC cultured in DMEM supplemented with different concentrations of Mg^2+^ **(e)** or in conditional medium from macrophages stimulated with different concentrations of Mg^2+^ **(f)**. **(g)** Representative TEM images and corresponding SAED patterns showing the precipitation formed in culture medium supplemented with different concentrations of Mg^2+^ scale bars = 50 nm. **(h)** XRD patterns of the precipitation formed in culture medium supplemented with different concentrations of Mg^2+^, stoichiometric HAp pattern (JCPDS card #09-0432) was shown as a reference. **(i)** The proliferation of MSC cultured in DMEM supplemented with different concentrations of IL-8.

**Table S1.**
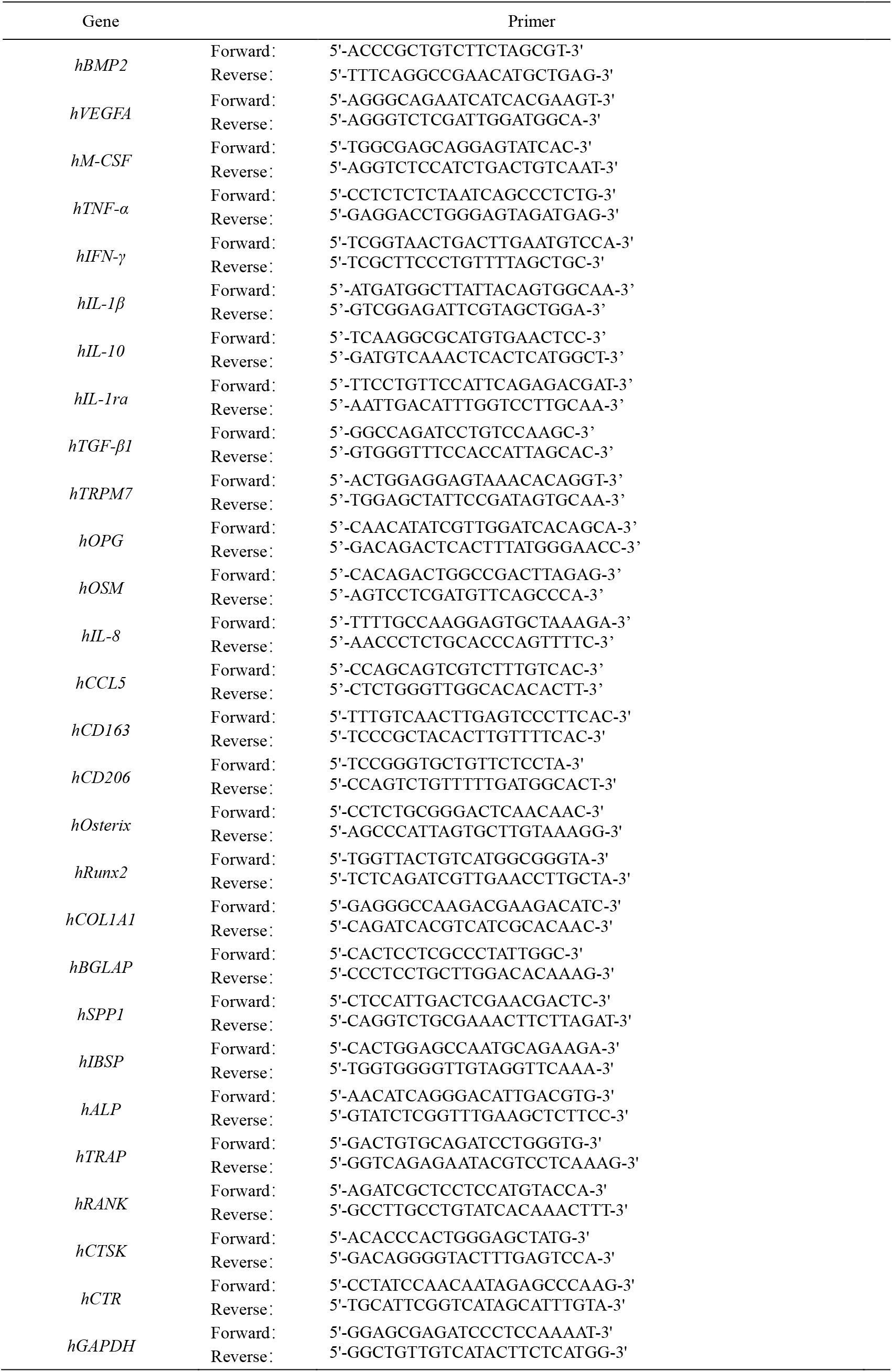
Primers used in the RT-qPCR assays.

